# AIControl: Replacing matched control experiments with machine learning improves ChIP-seq peak identification

**DOI:** 10.1101/278762

**Authors:** Nao Hiranuma, Scott M. Lundberg, Su-In Lee

## Abstract

ChIP-seq is a technique to determine binding locations of transcription factors, which remains a central challenge in molecular biology. Current practice is to use a “control” dataset to remove background signals from a immunoprecipitation (IP) target dataset. We introduce the *AlControl* framework, which eliminates the need to obtain a control dataset and instead identifies binding peaks by estimating the distributions of background signals from many publicly available control ChIP-seq datasets. We thereby avoid the cost of running control experiments while simultaneously increasing the accuracy of binding location identification. Specifically, AIControl can (1) estimate background signals at fine resolution, (2) systematically weigh the most appropriate control datasets in a data-driven way, (3) capture sources of potential biases that may be missed by one control dataset, and (4) remove the need for costly and time-consuming control experiments. We applied AIControl to 410 IP datasets in the ENCODE ChIP-seq database, using 440 control datasets from 107 cell types to impute background signal. Without using matched control datasets, AIControl identified peaks that were more enriched for putative binding sites than those identified by other popular peak callers that used a matched control dataset. We also demonstrated that our framework identifies binding sites that recover documented protein interactions more accurately.

## Introduction

Chromatin immunoprecipitation followed by sequencing (ChIP-seq) is one of the most widely used methods to identify regulatory factor binding sites and analyze regulators’ functions. ChIP-seq identifies the positions of DNA-protein interactions across the genome for a regulatory protein of interest by cross-linking protein molecules to DNA strands and measuring the locations of DNA fragment enrichment associated with the protein [3, 16, 30]. The putative binding sites can then be used in downstream analysis [39, 13], for example, to infer interactions among transcription factors [28, 34, 6], to semi-automatically annotate genomic regions [12, 15], or to identify regulatory patterns that give rise to certain diseases such as cancer [5, 4].

Identifying protein binding sites from signal enrichment data, a process called “peak calling,” is central to every ChIP-seq analysis, and has thus been a focus of the computational biology research community [47, 23, 48, 10, 19, 14, 33, 36, 21]. Like other biological assays, ENCODE ChIP-seq guidelines recommend that researchers obtain two ChIP-seq datasets to help separate desirable signals from undesirable biases: (1) an *IP (immunoprecipitation) target dataset* to capture the actual protein binding signals using immunopreciptation, and (2) a *control dataset* to capture many potential biases [24]. Peak calling algorithms compare IP and control datasets, locate peaks likely associated with true protein binding signals, and simultaneously minimize false positives. However, despite the guideline’s recommendations, many ChIP-seq users perform experiments either without a matched control dataset or with a related control dataset from a public database in order to avoid the additional time and expense of generating control datasets.

Here, we introduce AIControl (Figure 1(a)), a single-dataset peak calling framework that replaces a control dataset with machine learning by inferring background signals from publicly available control datasets on a large scale. As noted, most popular peak callers perform comparative ChlP-seq analysis using two datasets, – IP and control datasets. Many of them have an option to perform single-dataset analysis (i.e., IP dataset only) by determining the structure of background signals from the IP dataset itself; however, it is unlikely to be as accurate as when a control dataset is used. AIControl aims to estimate and simulate the true background distributions at each genomic position based on the weighted contribution of a large number of publicly available control datasets, where weights are learned from both the IP dataset and publicly available control datasets (see Methods for details).

**Figure 1.**
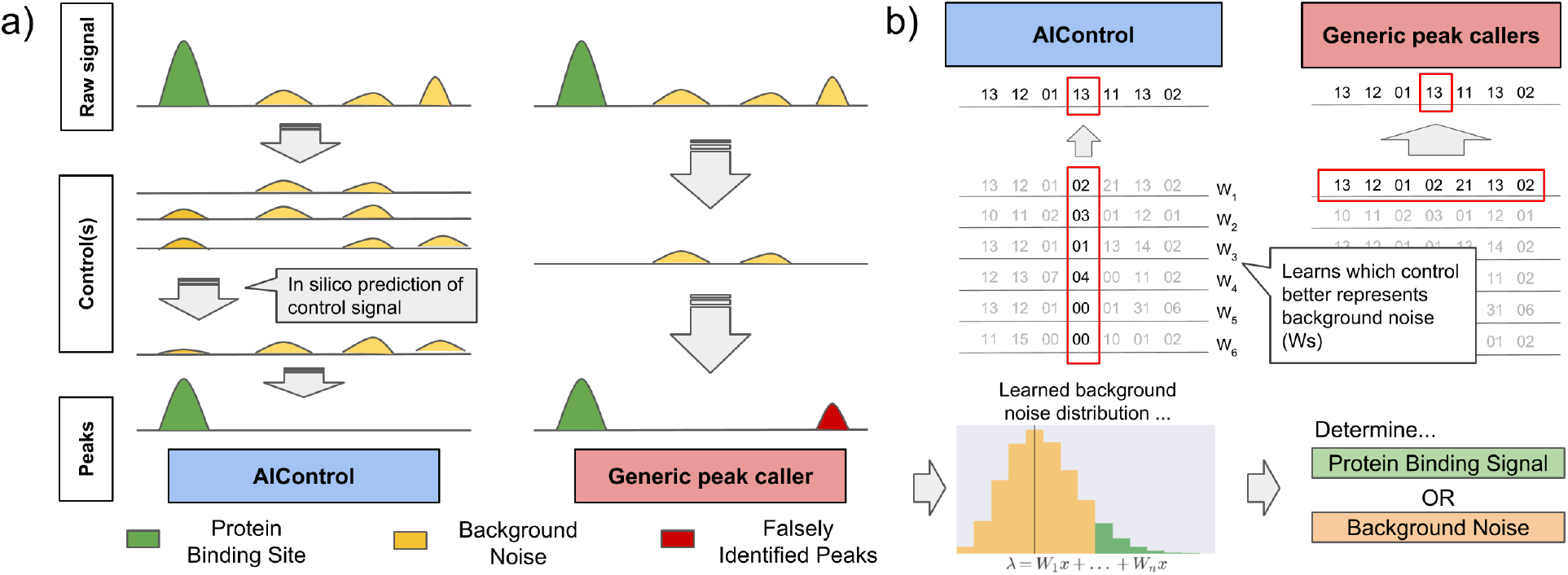
(a) An overview of the AIControl approach. A single control dataset may not capture different kinds of biases that give rise to background signal. AIControl more completely removes background signal of ChIP-seq by using a large number of publicly available control ChIP-seq datasets (see Methods). **(b) Comparison of AIControl to other peak calling algorithms**. (left) AIControl learns appropriate combinations of publicly available control ChIP-seq datasets to impute background signal distributions at a fine scale. (right) Other peak calling algorithms use only one control dataset, so they must use a broader region (typically within 5,000~10,000 bps) to estimate background distributions. (bottom) The learned fine scale Poisson (background) distributions are then used to identify binding activities across the genome.

Most popular peak callers – such as Model-based Analysis of ChIP-seq ver 2.0 (MACS2) [48] and Site Identification from Short Sequence Reads (SISSRs) [33] – learn local distributions of read counts from a matched control dataset in a nearby region (Figure 1(b), right). They then identify peaks by comparing observed read counts in the IP dataset to learned local background distributions across the genome. Several methods – such as MOdel-based one and two Sample Analysis and inference for ChIP-Seq Data (MOSAiCs) and BIDCHIPS – use a few predictors expected to represent sources of biases, such as GC content and read mappability. MOSAiCs performs negative binomial regression of an IP dataset using GC content, read mappability, and a matched control dataset as predictors [21]. Similarly, BIDCHIPS uses staged linear regression to combine GC content, read mappability, DNase 1 hypersensitivity sites, an input control dataset, and a mock control dataset [36].

AIControl’s main innovations are four-fold: (1) AIControl can learn position-specific background distributions at a finer resolution than traditional approaches by leveraging multiple weighted control datasets. Most other peak callers take a large window of nearby regions to learn the position-wise distributions, which may inaccurately estimate local structure of background signal. This feature also offers significant improvements over our previous work, CloudControl [14]. CloudControl generates one synthetic control dataset based on publicly available control datasets and uses peak callers to identify peaks that rely on a large window of nearby signals for background estimation. Throughout this paper, we show that AIControl significantly improves peak calling quality relative to CloudControl. (2) Existing peak callers require users to decide which control datasets to include. AIControl offers a systematic way to integrate a large number of publicly available control datasets. (3) Because AIControl integrates many control datasets, it can potentially capture more sources of biases compared to existing methods that use only one control (e.g., MACS2 and SISSRs). Most confounders – such as GC content and mappability – are likely present in some of the control datasets AIControl incorporates. See “Modeling background signal” in Methods for our mathematical formulation. (4) AIControl does not need a matched control dataset. We incorporate 440 control ChIP-seq datasets from 107 cell types in the ENCODE database. By inferring local structure of background signal from the large amount of publicly available data, AIControl can identify peaks even in cell types without any previously measured control datasets. We demonstrate that our framework intelligently uses existing control datasets to estimate background distributions for IP datasets in brand new cell types in a cross-cell-type setup.

We evaluated the AIControl framework on 410 ChIP-seq “IP datasets” available in the ENCODE database [8] (Table S1) using 440 ChIP-seq “control datasets” (Table S2 & S3). The IP datasets span across five cell types: K562, GM12878, HepG2, HeLa-S3, and HUVEC. Every cell type except for HUVEC is a cell line, and HUVEC is a primary cell (endothelial cell of umbilical vein). Results show the following. (1) AIControl outperformed other peak callers on identifying putative protein binding sites based on sequence-based motifs (Figure 2). All competing peak callers used matching pairs of IP/control datasets, whereas AIControl did not – it used only IP datasets (no matching control) and publicly available control datasets. AIControl predicted putative binding sites well even when all control datasets from the same cell type were removed, which suggests that it reliably estimates background signals in a cross-cell-type manner when ChIP-seq is performed on a new cell type (Figure 4). (2) PPIs were more accurately predicted from peaks called by AIControl than from those called by all other methods in all tested cell types (Figure 5). (3) Peaks identified by AIControl showed superior performance in motif enrichment and PPI recovery tasks when they were processed with the irreproducible discovery rate (IDR) pipeline (Figure S1). (4) AIControl exhibited strong performance on datasets that are not part of the ENCODE database (Figure 6). Our findings suggest that AIControl can remove the time and cost of running control experiments while simultaneously identifying binding site locations of transcription factors accurately.

**Figure 2.**
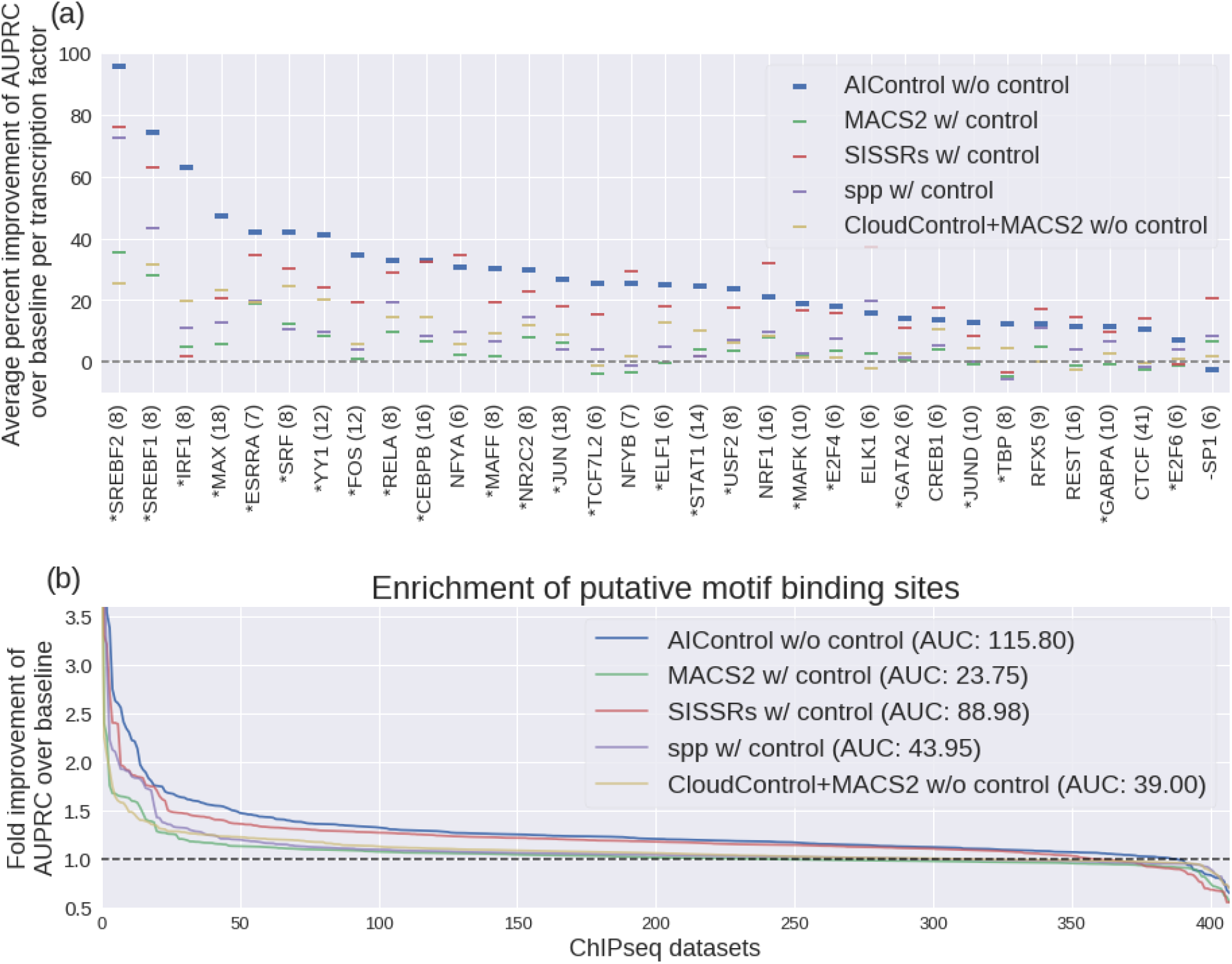
(a) Average percent improvement of area under precision recall curves (AUPRCs) averaged per transcription factor. Peaks were identified using: (1) AIControl w/o control, (2) MACS2 w/control, (3) SISSRSs w/control, (4) SPP w/control, and (5) CloudControl + MACS2 w/o control. Only the transcription factors that were measured more than 5 times are shown (the number of measurements shown in parenthesis), and they are ordered by AIControl’s performance. The transcription factors that AIControl performed the best on are shown with an asterisk (24 out of 33). The ones on which AIControl performed worst are shown with a minus sign (1 out of 33). **(b) Relative performance of five peak calling methods compared to MACS2 without using a matched control dataset as a baseline (dotted line)**. The y-axis shows the fold improvement of the area under the precision-recall curves (AUPRCs) for predicting the presence of putative binding sites with significance values associated with the peaks over the baseline (i.e., MACS2 without using a matched control dataset) across the whole genome. The x-axis shows the all 410 ENCODE ChIP IP datasets ordered by the fold improvement (y-axis) across all tested cell type; 149, 99, 60, 87 and 15 for K562, GM12878, HeLa-S3, HepG2, and HUVEC, respectively. Note that the ordering of datasets is different for each peak caller. Area between each line and the dotted baseline is shown in parenthesis.

## Material & Methods

### Methods

#### Modeling background signal

AIControl models background signals across the human genome as a linear combination of multiple different sources of confounding biases. In particular, let us denote a control ChIP-seq dataset *i* as *y_i_* ∈ ℝ^*g*^, where *g* represents the number of binned regions across the whole genome. Let us also denote the signals from *n* bias sources as *x*_1_,…,*x_n_* ∈ *R^g^*. For example, *x*_1_ may represent the GC content across the whole genome. Then, we model each control dataset *y_i_* as a linear combination of *x*_1_,…, *x_n_*:

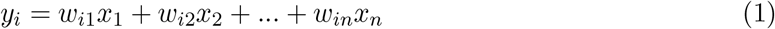

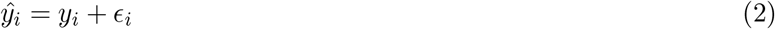

Here, *ϵ_i_* represents irreproducible noise in a control dataset *i*, and *ŷ_i_* represents an observed control dataset *i*. Each control dataset is modeled as a specific linear combination of *n* bias sources, and *w_i_* = (*w*_*i*1_,*w*_*i*2_,…,*w_in_*) ∈ ℝ^*n*^ corresponds to a specific control dataset i. These weight vectors of all control datasets are *not* observed.

For a particular target IP dataset *t*, AIControl attempts to estimate its background signal *ŷ_t_*, which is modeled as a weighted linear combination of *x*_1_,…,*x_n_* with a weight vector *ŵ_t_* ∈ ℝ^*n*^:

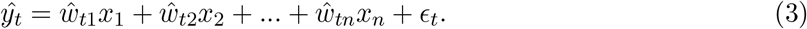

Below, we show that we can estimate *ŷ_t_* without explicitly learning *ŵ_t_* and *x*_1_,…,*x_n_*. The idea is that we can view a set of weight vectors *w*_1_,…*w_m_* ∈ ℝ^*n*^ from m publicly available control datasets (here, 440 ENCODE control datasets, summarized in Table S2 and S3) as a spanning set of ℝ^*n*^ (or a large subset of it) provided that *n* <<*m*. Thus, we can model *ŵ_t_* as a linear combination of weight vectors *w*_1_,…*w_m_*:

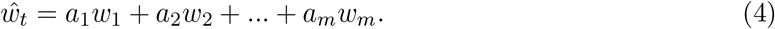

Plugging equation (4) into equation (3) leads to:

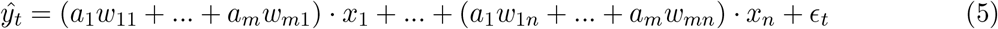

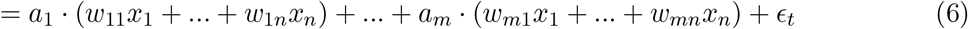

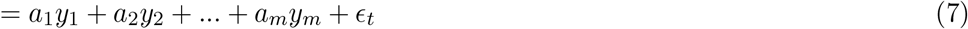

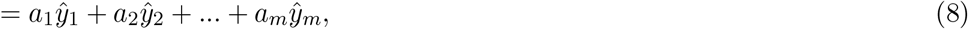

where *ϵ_t_* represents the total irreproducible noise. This shows that *ŷ_t_* can be represented as a weighted linear combination of a large number of *m* control datasets. To learn the coefficient vector, *a* = 〈*a*_1_,…,*a_m_*〉, we could do a linear regression of a true background-signal vector for IP dataset *t, ŷ_t_*, against *ŷ*_1_,…,*ŷ_m_*; however, *y_t_* is not observed. Instead, we regress the observed signal of the IP dataset, *o_t_*, against *ŷ*_1_,…,*ŷ_m_* given that *o_t_* can be decomposed as follows.

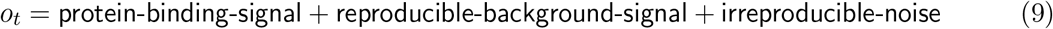

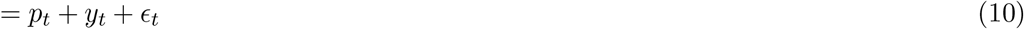

The idea is that in theory, *m* control datasets, *ŷ*_1_,…,*ŷ_m_*, should contain no information about *p_t_* and *e_t_*; therefore, we can determine the coefficient vector *a* by regressing *o_t_* against *ŷ*_1_,…,*ŷ_m_* unless we overfit. Here, the sample size is millions, and the number of variables is 440, which means that this problem is far from high-dimensional and unlikely to overfit.

#### Computing coefficients

We regularize AIControl by applying the L2 ridge penalty on the coefficient vector *a* = 〈*a*_1_,…, *a_m_*〉. This leads to the following objective function:

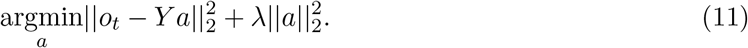

Here, *Y* is a *g* by *m* (= 440) matrix, where each column *i* corresponds to *ŷ_i_*, the ith observed control dataset. Using the closed form solution of ridge regression, we can efficiently compute the coefficient vector, *a*:

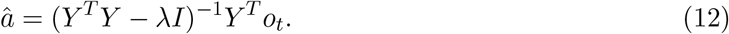

Because this regression problem involves a large number of samples (i.e., is far from being high-dimensional), we chose a small regularization coefficient λ = 0.00001 to ensure numerical stability. Since the dimension of *Y* is *g* by *m*, where *m* is 440 and g is 30 million (under the default setting where the size of bins is 100 base pairs (bps)), we are unlikely to require strict regularization to prevent overfitting.

When implementing AIControl, we learned separate models for signals mapped to forward/reverse strands and even/odd positions, which results in four coefficient vectors *â* per target IP dataset. `reproducible-background-signal` is estimated separately for forward and reverse strands as *ŷ* _forward_ and *ŷ*_reverse_ by applying the coefficients learned at even positions to calculate *ŷ* at odd positions, and vice versa. Training on odd positions and predicting on even positions (and vice versa) is designed to further prevent any possible overfitting. Spearman’s correlation values of learned coefficients are shown in Figure S2. These values are generally above 0.8 for any pair in the same IP dataset, showing that learned sets of weights are consistent among forward, reverse, odd- and even-positioned data.

It is important to note that we need not to recompute *Y^T^Y* ∈ℝ^440×440^ for different IP datasets because it remains constant when the same set of control datasets is reused. To estimate *ŷ_t_*, we need only two passes through the whole genome: the first to compute *Y^T^ot*, and the second to calculate *âY*.

#### Identifying peaks

Commonly used peak calling approaches identify a peak based on how far its read count at a particular genomic region diverges from the null distribution (typically, Poisson, Zero-inflated Poisson, or negative binomial distribution) that models background signal without protein-binding events [48, 33]. Usually, null distributions are semi-locally fit to signals from nearby regions (5000~10000 bps) in a matched control dataset.

Like many other peak callers, AIControl uses the Poisson distribution to identify peak locations; however, null background distributions are learned at much finer scale. In particular, we use the following probabilistic model of the null background distribution for the read count observed at the *i*th position of genome *c_ti_* in the target IP dataset *t*:

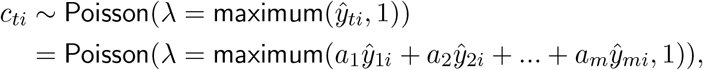

where *ŷ*_1_,…,*ŷ_m_* represent *m* publicly available control datasets, and *a*_1_,…, *a_m_* are estimated using equation (12). This approach can be viewed as fitting a Poisson distribution to count data at each genomic bin *i, ŷ*_1*i*_,…, *ŷ_mi_*, that are weighted differently with corresponding weights *a*_1_,…, *a_m_*. The use of *m* control datasets (not just one matched control) lets us learn a higher resolution background distribution (Figure 1(b)). Finally, we introduce the minimum base count of 1 read to prevent *ŷ_t_* from being too small or negative since the coefficient vector *a* can contain negative values. In our implementation, users have an option to include nearby *b* bins to learn the null background distribution in case they choose not to use our standard control dataset release and so do not have a sufficiently large number of background controls *m*.

We then calculate the p-value and fold enrichment of the observed count at each genomic bin based on the learned null background distribution and background count. To this point, peak identification processes are completed separately for forward and reverse strands; we use *a*_1_,…,*a_m_* learned from even-numbered regions to identify peaks at odd-numbered regions (and vice versa) for each forward and reverse strand. We then slide the locations of the p-values and enrichment values by 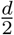 and 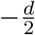, for forward and reverse signals respectively. *d* is defined as the expected distance between forward and reverse peaks; it is automatically estimated in our framework. Finally, the smaller negative log10 p-value and fold enrichment of read counts between the slid forward and reverse signals at every position is output as a peak signal. This last step ensures that peaks have bimodal shapes as expected for transcription binding signals [48].

#### Estimating distance between forward and reverse peaks *d*

AIControl automatically estimates the distance between forward and reverse peaks similar to other peak callers. Specifically, for each dataset, we find the sliding distance *d* that minimizes the disagreement between the forward and reverse mapped reads. In particular, the disagreement is defined as follows:

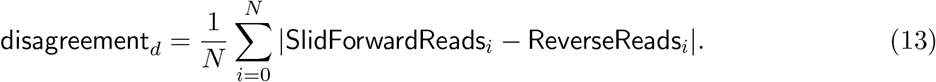

*N* is the number of bins in the hg38 genome, which is approximately 30 million with a bin size of 100 bps. We find d that minimizes the disagreement value with brute force search between *d* = 0 to *d* = 400. With a default binning size of 100 bps, there are only four options for *d*, and it is relatively fast to find the optimal *d*. The summary of *d* estimation for all 410 tested ENCODE IP datasets is shown in Figure S3.

#### Merging contiguous bins with significant binding signal

AIControl assigns a p-value and fold enrichment of binding signal to each 100 bp genomic bin. As an optional post-processing step, the current implementation of AIControl can merge contiguous bins that have more significant binding signal than threshold (default is negative log10 p-value of 1.5, approximately p-value of 0.03) by taking the maximum p-value and fold enrichment values among them. The resulting peaks are output in a .narrowPeak format.

### Data processing

#### Aligning BAM files

We describe BAM files used in this project in our prior work on ChromNet [28]. Specifically, the raw FASTQ files were downloaded from the ENCODE database and were mapped to the UCSC hg38 genome with BOWTIE2 to ensure an uniform processing pipeline [25]. We provide the full list of ENCODE experimental IDs used in this project in Supplementary Data 1.

#### Calling peaks with other methods

The version of MACS2 used in this paper was MACS2 2.1.0.20150731 [48]. The peaks were called with the following command: “`macs2 callpeak -f BAM -t chipseq_dataset -control matched_control -q 0.05`”. The version of SPP used was 1.13, and the peaks were called with an FDR threshold of 0.05 using the “`find.binding.positions()`” function in its R package [19]. Additionally, we downloaded SPP peaks from the ENCODE portal if they were available (we call it “SPP-ENCODE”). For SISSRs, we used v1.4, and the peaks were also called with a p-value threshold of 0.05 with the following command: “`sissrs.pl -i chipseq_dataset.bed -b matched_control.bed -p 0.05 -s 3209286105`” [33]. The peaks from CloudControl were obtained in conjunction with MACS2 using the same parameters as above [14]. All resulting peak files can be viewed in Google Drive through our GitHub repository under the “Paper” section (https://github.com/hiranumn/AIControl.jl).

#### Obtaining peaks that are optimally controlled with IDR

The ENCODE official pipeline for processing biological replicate samples is to use SPP and IDR in combination [26]. We also investigated the performance of peak callers in combination with the IDR process (Figure S1). In particular, for peaks processed with SPP, we downloaded peak files tagged as “optimal idr thresholded peak” from the portal website of ENCODE [8]. If they were available on the hg19 genome, we used the UCSC liftover tool to convert peak locations from the hg19 to hg38 genome. For other peak callers (i.e., MACS2, SISSRs, and AIControl), we used the Python implementation of idr (https://github.com/kundajelab/idr) to re-order peaks for each pair of biological replicates. We believe that the significant value thresholds we used (0.05 for MACS2 and SISSRs, and 0.03 for AIControl) are lenient enough to capture both reproducible and irreproducible signals that are required for the IDR process.

#### Storing large matrix of control signals efficiently

One of the challenges in implementing AIControl in a user-friendly manner is to find an efficient way of storing a massive amount control datasets. In particular, we have 440 ChIP-seq control datasets, and each of them is represented as a 30 million long vector, which stores read count for every 100 bp bin. Collectively, the control datasets are represented as a sparse, non-negative matrix of size 440 by 30 million. For this project, we developed our own file format to store the large matrix. First three 8 bit chunks of the file encode the following three parameters: (1) a number of control datasets (i.e. width of the matrix), (2) a maximum possible value stored in the matrix (100, if duplicate reads are removed), and (3) a data type (i.e., UInt8 or UInt16). Subsequent 8 or 16 bits, depending on the data type, are used for indicating an actual value in the current entry of matrix, if it is less than the predefined maximum value. Otherwise, it is used for indicating how many entries (value – the maximum value) to skip column-wise by filling in 0s. With this file format, we can compress all 440 control datasets to 4.6 GB. Given that typical BAM files are about 0.5~3.0 GB, we believe this makes our standard control dataset package compact enough for users to download.

### Analysis pipeline

#### Standardizing peak signals

Different peak calling algorithms identify different numbers of peaks at a given significance value threshold and generate peaks with different widths. To eliminate the possibility that these differences in peak numbers or widths create biases, we standardized peak signals for each dataset as follows. (1) We bin peak signals by 1,000 bp windows. This creates vectors where each entry corresponds to the peak with the largest ranking measure (i.e., negative log10 p-values or signal values) among the peaks that fall into the corresponding bin. We thereby standardize peak width to 1,000 bps across all methods. (2) For each dataset, we use the top *n* peaks for all peak callers, where *n* is the minimum number of peaks identified from all tested peak callers at a p-value cutoff of 0.05. Choice of ranking measure matters. We used column 7 (signal value) of the narrowPeak format for SPP and AIControl, and column 8 (p-value) for MACS2. For SISSRs, its output does not follow the narrowPeak format, but we use p-values associated with peaks as the ranking measure. This process, which standardizes the number of peaks identified by different peak callers, results in an average of 21,470 genome-wide peaks per dataset (Figure S4).

#### Evaluating motif enrichment

We applied AIControl to 410 ChIP-seq IP datasets from the ENCODE database for which we could find motif information. For each IP dataset, we obtained a probability weight matrix (PWM) of binding sites for its target transcription factor from the JASPAR database [18]. We then used FIMO from the MEME software to search for the putative binding sites at the p-value threshold of 10^-5^ [2]. The idea is that correctly identified peaks are likely in a region that contains the corresponding motif. Of course, motif enrichment alone may not be a reliable measure; thus in addition to examining the whole genome, we also focus on the regions where transcription factor binding occurs relatively more often. For instance, studies show that 98.5% of the transcription factor binding sites are positioned in DNase 1 hypersensitivity (DHS) regions [44].

Therefore, to increase the reliability of motif-based evaluation criteria, we focused our analysis on the following four regions when we performed motif enrichment-based evaluation: (1) the whole genome, (2) DNase 1 hypersensitivity regions, (3) regions that are 5000 up and downstream of the start sites of protein coding genes, and (4) regions that have more than 50% GC content. The DHS signals were downloaded from the portal website of the Roadmap Epigenomics project [22]. The regions proximal to protein coding genes were obtained through BioMart [20]. After standardizing peak signals across all methods (as described in the previous subsection), for each peak calling method, we predicted the presence of putative binding sites in each of aforementioned regions using varying thresholds of the significance of peaks. This led to a standard precision-recall curve for predicting the presence of putative binding sites when the significance level of the peak varies (i.e., x-axis in the standard precision-recall curve). We then used the area under the precision-recall curve (AUPRC) to assess peak calling method performance. We computed the AUPRC using Riemann sum approximation.

We used “waterfall plots” to collectively visualize the AUPRCs of all peak callers for all IP datasets in each cell type for (1) the whole genome (Figure 2 and S5), (2) DHS regions (Figure S6), (3) up-/down-stream regions of protein-coding genes (Figure S7), and (4) high GC content regions (Figure S8). For example, in Figures 2 (b) and 4, each colored line corresponds to the performance of a particular peak calling algorithm on IP datasets. The y-axis measures the ratio of AUPRC given by the corresponding peak calling methods to AUPRC given by the baseline method, i.e., MACS2 without a control dataset (also represented by the dotted line). The x-axis corresponds to the IP datasets, which are sorted independently for different peak calling methods based on their y-axis values. We removed datapoints if any peak caller called less than 20 peaks to ensure the stability of AUPRC values (Figure S4 (b)). Peak callers with larger areas under the colored line and above the dotted line relative have, on average, better identified peaks supported by sequence motifs (Table S5 and S9). The AUPRC value for each peak caller on each dataset is shown in Supplementary Data 2 for the whole genome, Supplementary Data 3 for the DHS regions, Supplementary Data 4 for the gene proximal regions, and Supplementary Data 5 for the GC rich regions.

#### Using area under PR curve of *n* most significant peaks as an evaluation metric

As described in ‘standardizing peak signals’ above, we analyze only the *n* most significant peaks, where *n* is determined by the minimum number of peaks called among all peak callers. We then generated the precision-recall curve for predicting the presence of putative binding sites using peak significance values and used the AUPRC to assess peak calling quality. Here, we aim to justify the use of AUPRC as an evaluation metric. Figure S9 explains what the AUPRC metric captures for peak callers with different behaviors. Figure S9 (a) shows a precision-recall curve when a peak caller performs well at selecting true peaks in top *n* (captured by area A) but poorly at ordering them in top *n* (captured by area B). Figure S9 (b) shows an example for the opposite case, in which a peak caller perform poorly at placing true peaks in top *n* but well at ordering them in top n. Both quantities measured by the area A and B are important for high quality peak calling. In practice, researchers use 500 ~ 10000 most significant peaks depending on transcription factors. Our average choice for *n* is 21,470, much higher than the widely used threshold (Figure S4 (a)).

#### Obtaining and evaluating on protein-protein interaction (PPI) matrix

The validated PPI interactions we used for evaluation were downloaded from the BioGrid website by 2018/2/5 [42]. We used only the PPIs in Homo sapiens from `BIOGRID-ORGANISM-Homo_sapiens-3.4.157.mitab.txt`. Because the interactions are recorded in terms of Entrez ID in BioGrid, the uniprot IDs of the targeted transcription factor of ENCODE IP datasets were converted to Entrez ID using the Uniprot Mapping Tool from http://www.uniprot.org/mapping/.

PPIs were estimated for each cell type as follows. First, for each peak calling method, the inverse correlation matrix from all *n* IP datasets in the cell type of interest was computed using standardized peak signals (see ‘Standardizing peak signals’) after binarization. This resulted in a matrix of size *n* by *n*. Finally, the magnitudes of the inverse correlation values were used as predictors for PPIs.

To visualize the quality of predictions, we used fold enrichment plots (Figures 5 and S10), like we did previously [28]. Fold enrichment is defined as follows for given number of selected predicted interactions (x-axis of Figures 5 and S10):

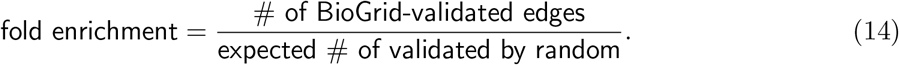

This value has been shown to reflect both type 1 and type 2 errors. We plot the fold enrichment value (y-axis) against the number of predicted interactions selected (x-axis) (Figures 5 and S10). A larger area under the fold enrichment curve indicates superior performance similar to PR curves.

#### Measuring consistency among pairs of unrelated IP datasets

This analysis used 9,310 pairs of IP datasets in K562 that target unrelated transcription factors. Here, “unrelated” means that the pair of transcription factors has no documented PPIs based on the BioGrid database [42]. The number of shared peaks between a pair of datasets is computed as follows: First, we binarize the standardized peak signals for each dataset. Then, we counted the number of non-zero entries at the intersection between two datasets. This gives us the number of peaks in the same genomic locations between a pair of datasets.

### Software availability

The Julia 1.0 implementation of the AIControl software and a thorough step-by-step guideline can be found at our github repository: https://github.com/hiranumn/AIControl.

#### Aligning an input .fastq file to the hg38 genome

Users must align their input .fastq files to the hg38 genome from the UCSC repository, which can be found at http://hgdownload.soe.ucsc.edu/goldenPath/hg38/bigZips/hg38.fa.gz using bowtie2 [25]. Unlike other peaking callers, the unique core idea of AIControl is to leverage all available control datasets in public. This requires all data (both your target ChIP-seq and public control datasets) to be mapped to the exact same reference genome. Currently, the control datasets that we provide are mapped to the UCSC hg38 genome. Therefore, for instance, if an input ChIP-seq data is mapped to a slightly different version of the hg38 genome, the AIControl pipeline will report an error. If you start with a `.bam` file that is already mapped, the recommended way of resolving this error is to use `bedtools`, which provides a way to convert `.bam` files back to `.fastq` files and realign to the correct version of the hg38. (See “Step 1” and “Step 3.1” on our github repository at https://github.com/hiranumn/AIControl for specific commands).

#### Converting and sorting a .sam file

The direct output `bowtie2` is a `.sam` file. Users need to use `samtools` to convert it to a `.bam` file and sort it in lexcographical order (See “Step 2” and “Step 3” on our github repository at https://github.com/hiranumn/AIControl for specific commands).

#### Downloading compressed control data files

AIControl requires users to download binned control datasets on their local systems. The compressed control data files for all 440 ENCODE control datasets are available through Google Drive on the GitHub page under the “Paper” section or our project website at https://sites.google.com/cs.washington.edu/suinlee/project-websites/aicontrol under the “Data” section. We have two separate files for forwardly and reversely mapped signals. Each one is 4.6GB and 13GB when unfolded (See “Step 4” on our github repository at https://github.com/hiranumn/AIControl).

#### Running the AIControl script

We generated a julia script file, `aicontrolScript.jl`, that performs the AIControl framework, which takes in the sorted `.bam` file and outputs a `.narrowPeak` file (See “Step 5” on our github repository at https://github.com/hiranumn/AIControl for how exactly to execute it). We have tested and validated our pipeline on Ubuntu 18.04, MacOS Sierra, and Windows 8.0. Although AIControl works on all tested systems, we recommend that users run AIControl on Unix based systems (e.g. macOS or Ubuntu), because the other peripheral software, such as `samtools` or `bowtie2` are easier to install there. Please refer to the “Issues” page of our github repository at https://github.com/hiranumn/AIControl or, if you have a problem running the AIControl pipeline.

## Results

### Peaks identified by AIControl are more enriched for binding sequence motifs

We compared AIControl to the following four alternative peak calling methods in terms of its enrichment for putative binding sites, the most widely used evaluation metric for peak-calling algorithms: MACS2 [48], SISSRs [33], SPP [19], and MACS2 + CloudControl [14]. To define putative binding sites without using ChIP-seq data, we identified sequence motifs using FIMO from the MEME tool [2] and position weight matrices (PWMs) from the JASPAR database [18] (see Methods). MACS2, in particular, has been favored by the research community due to its simplicity and steady performance as validated by many comparative studies of peak calling algorithms [43, 47, 23]. To evaluate the enrichment for putative binding sites, we used ranking measures (negative log10 p-values or signal values, see “Standardizing peak signals”) of peaks to predict the presence of putative binding sites and measured the area under the precision-recall curves (AUPRCs) in the following four genomic regions: (1) the whole genome, (2) DNase 1 hypersensitivity regions (DHS), (3) 5 kilo-base up and downstream of protein coding gene start sites, 4) regions with more than 50% GC content. To ensure that each peak caller was tested on the same number of peaks, we measured AUPRC values on the *n* most significant peaks, where *n* is the minimum number of peaks called across all peak callers for each IP dataset (see Methods). This process prevents peak callers that identify more peaks at a given threshold from having an unfair advantage. For the analyses across the whole genome, this resulted in an average *n* of 21,470 peaks per IP dataset for the entire genome (Figure S4).

Figure 2 (b) compares the AUPRCs across the whole genome achieved by the five peak callers for 410 IP datasets across five different cell types: K562 (149), GM12878 (99), HepG2 (87), HeLa-S3 (60), and HUVEC (15). AIControl yielded better fold improvements of AUPRCs over that of baseline than the other peak callers (p-value<0.0001 with the Wilcoxon signed-rank test on AIControl vs SISSRs on matched pairs of fold improvements). When the results are viewed separately for the five cell types, AIControl achieves the best performance in all cell types except for HUVEC (Figure S5 and Table S5). Again, AIControl used only IP datasets without their matched control datasets, whereas other peak callers, except for CloudControl+MACS2, accessed both IP and their matched control datasets. AIControl continues to perform better on the motif enrichment task even when the analyses are restricted the aforementioned regions (2)-(4) of the genome. (Table S6, S7, S8 and Figure S6, S7, S8). To validate that we used these peak callers correctly, we further investigated the performance of all peak callers on five IP datasets for the RE1-Silencing Transcription factor (REST) measured in K562, for which we had quantitative polymerase chain reaction (qPCR) verified TF binding sites [32]. All five peak callers identified all eight qPCR-confirmed binding locations on chromosome 1.

Datasets that target certain transcription factors yielded more performance improvements than others. Figure 2 (a) shows the mean percent improvement of the five peak callers – three of which use matched control datasets – over base (i.e. MACS2 without matched control) for transcription factors that are measured more than five times across all 410 IP datasets. The AIControl framework outperformed all other peak callers in 24 out of 33 transcription factors without needing a matched control experiment. In particular, our framework exhibited major average improvements on transcription factors MAX, JUN and STAT1, over the best performing peak callers, while it showed decreased performance on SP1. Arvey et al. (2012) [1] examined cell-type-specific binding of patterns of 15 transcription factors from ENCODE data between K562 and GM12878. In the study they observed more cell-type-specific peaks from transcription factors MAX, JUN, JUND, YY1, and SRF. In Figure 2 (a), AIControl performs the best on all of these factors, suggesting that our framework is able to well identify binding sites of transcription factors that exhibit differential binding patterns depending on target cell types.

We also investigated whether our analysis is affected by the quality of putative true binding sites by checking the relationship between the relative performance of AIControl over MACS2 across the whole genome and the information contents of the JASPAR PWMs. However, we did not find any significant correlation (r=0.02, p-value=0.84, Figure S11).

Interestingly, but not surprisingly, the more publicly available control datasets are incorporate, the better the performance of AIControl is. We picked 36 datasets that are associated with transcription factors measured more than 10 times across all tested cell types (i.e., YY1, CEBPB, STAT1, JUN, FOS, REST, MAX, CTCF, and NRF1 from each tested cell type; some TF/cell-type pairs were missing due to data availability). When averaged over many random subsets of different sizes, we observed that the AIControl framework experiences monotonic but diminishing increase of the performance from no control to 440 control sets, peaking at 29% improvement with all 440 control sets (Figure S12). While approximately 86% of the overall improvement is made with 200 control datasets, the additional 240 datasets still contribute positively (p-value<0.001 for Wilcoxon signed-rank test, comparing the performance at 200 and 440 control pairwise). This result suggests that the improved performance of AIControl in fact depends on the inclusion of public control datasets.

We recognize that our own pipeline for SPP is different compared to that of the ENCODE consortium since it has extra read/peak filtering steps. Naturally, this results in slightly different sets of peaks. Therefore, we decided to compare AIControl directly against the ENCODE peaks (SPP-ENCODE) in order to assure that AIControl still holds an advantage. We downloaded a peak file from the ENCODE portal for each dataset if it was available. Figure S13 shows that SPP-ENCODE outperforms SPP potentially due to the extra filtering steps, and more importantly, AIControl outperforms SPP-ENCODE. We note that we did not include SPP-ENCODE to Figure 2 because the SPP-ENCODE data is available in only a subset of IP datasets (i.e., 123 out of 149 for K562; 65 out of 99 for GM12878; 46 out of 87 for HepG2; 18 out of 60 for HeLa-S3; 7 out of 15 for HUVEC). The IDs of SPP-ENCODE peak files downloaded are listed in Supplementary Data 6.

### AIControl coefficients reflect cell type specificity but not lab specificity

AIControl learns the weights of contributions by all 440 ENCODE control datasets to estimate the background ChIP-seq signals for each IP dataset (Figure 1; also see Methods). Figure 3 shows the magnitude of weights assigned to all 440 control datasets (columns) for each of the 410 IP datasets (rows). A clear block diagonal pattern emerges when we sort the rows and columns based on cell type (Figure 3). This is expected because known factors for background signal signals, such as sonication bias and DNA acid isolation, depend on cell types. On the other hand, when we sort the rows and columns based on lab, we see less significant pattern, except for the datasets from the Weissman lab (Figure S14). This suggests that lab-specific batch effects are less significant than cell type-specific effects on the ENCODE ChIP-seq data. Although the control datasets from the same cell type as the IP dataset are more likely to have large weight magnitudes (Figure 3), it is important to note that AIControl learns to put high weights on some of the other biologically similar cell types. For example, the green box in Figure 3 indicates the weights of control datasets measured in GM12892 and learned for the IP datasets in GM12878. Both are B-lymphocyte cell types, and AIControl learns to leverage information from both cell types to identify peaks more accurately in GM12878.

**Figure 3.**
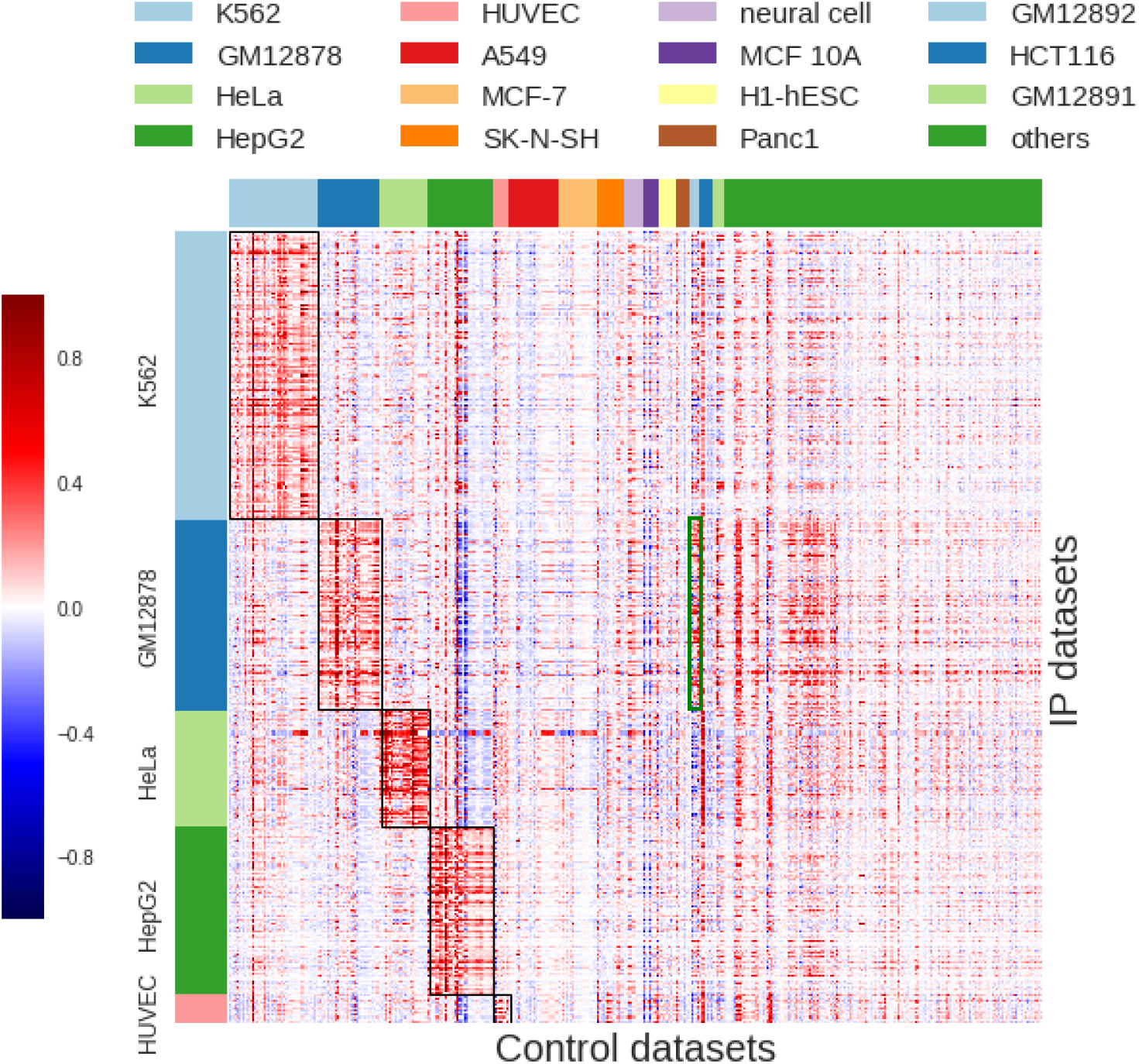
Normalized weights the AIControl model assigns to 440 ENCODE control datasets (columns) for each of the 410. IP datasets (rows) (Table S1 and S3). The black rectangles indicate the weights of the control datasets measured in the same cell type as the IP datasets. The green rectangle indicates the weights for control datasets measured in GM12892, which AIControl learned to estimate background ChIP-seq signals for IP datasets measured in GM12878. Both are B-lymphocyte cell types.

### AIControl performs well in a cross-cell-type setting where control datasets from the same cell types or the same labs are removed

The results described in the previous subsection indicate that AIControl leverages information about background signal from biologically similar cell types. A natural question is whether AIControl can correctly identify peaks in an IP dataset from a new cell type that is not included in the public control datasets that AIControl uses. This tests AIControl’s ability to estimate, in a crosscell-type manner, background signals in an unknown cell type from background signals in known cell types. Another important question is whether AIControl performs well for an IP dataset generated in a lab that did not generate the control datasets AIControl uses. To address these questions, we compared the following settings: (1) AIControl with all 440 control datasets except for matched controls, (2) AIControl without control datasets from the same cell type as the IP dataset, (3) AIControl without control datasets from the same lab as the IP dataset, (4) AIControl without control datasets from the same lab or the same cell type as the IP dataset, (5) SISSRs with matched control datasets, and (6) SISSRs without matched control datasets. We chose SISSRs because it is the best competitor in terms of identifying presence of motif sequences (Figure 2).

Figure 4 shows that AIControl with different patterns of excluding control datasets are still able to outperform SISSRs with matched control datasets in all cell types except for HUVEC, for which we only have a small number of datasets (n=15). Notably, the exclusion of control datasets from the same cell type (i.e., setting (2) and (4)) has a larger impact on the performance than the lab-based exclusion (i.e., setting(3)). This indicates that lab specific biases are less significant and easier to learn in a cross-lab setting. For setting (2) and (4), we observed the largest decrease in the performance of AIControl in K562, followed by HepG2 and HeLa-S3. On the other hand, in GM12878, AIControl was able to leverage information from other B-lymphocyte cell lines(i.e., GM12892) (Figure S15).

**Figure 4.**
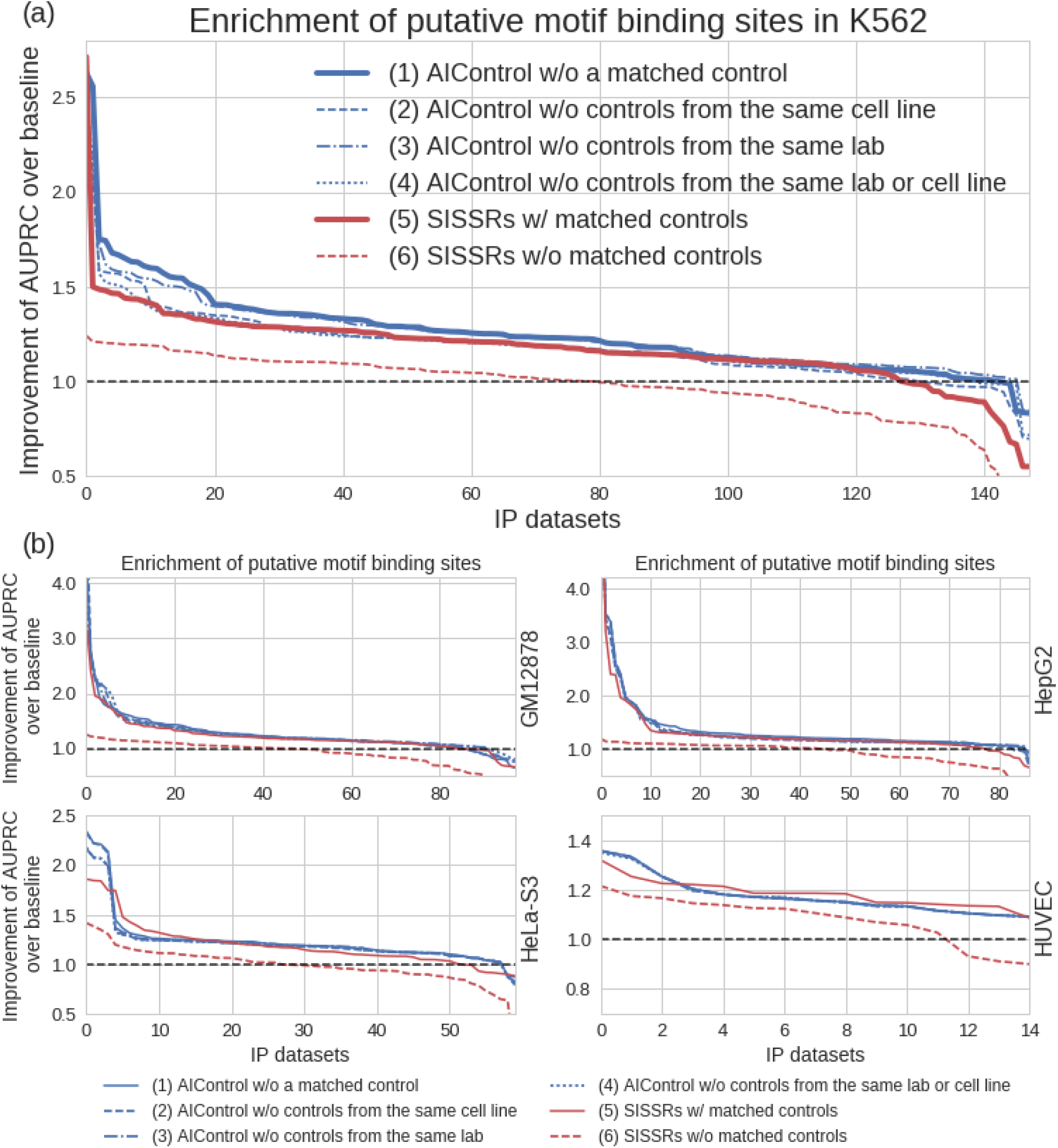
Relative performance of AIControl when control datasets from the same cell types or the same labs have been removed. We compare across six settings: (1) AIControl with all 440 control datasets except for matched controls, (2) AIControl without control datasets from the same cell type as the IP dataset, (3) AIControl without control datasets from the same lab as the IP dataset, (4) AIControl without control datasets from the same lab or the same cell type as the IP dataset, (5) SISSRs with matched control datasets, and (6) SISSRs without matched control datasets As in Figure 2, the y-axis shows the fold improvement of the AUPRCs for predicting the presence of putative binding sites compared with the baseline (i.e., MACS2 without using a matched control dataset) for (a) 149 IP datasets measured in K562, and (b) IP datasets measured in the tier 1 ENCODE cell types: GM12878, HepG2, HeLa-S3, HUVEC.

The decline in K562, followed by HepG2 and HeLa-S3, has two likely reasons. First, the largest number (53 of 440, or 12.0%) of ENCODE control datasets are from the K562 cell line (Table S3). Second, the structure of background signals in K562, HepG2, and HeLa-S3 cell types may be unique because of their abnormal karyotypes. Regions with multiplication, deletion, and copy number variations that are not documented in the reference genome can display signals that are locally proportional to alterations in the abnormal karyotype. This makes it harder for AIControl to estimate background signals in abnormal regions without having access to the controls from the same cell type. On the other hand, in GM12878 and HUVEC, which is known to have a normal karyotype, the performance of AIControl did not drop even without having access to control datasets from the same cell types.

We further investigated whether including additional control datasets with abnormal karyotypes hurts the performance of AIControl because they are less informative in certain regions, or including control datasets from just normal karyotypes is better even if it results in a significantly reduced number of datasets. For the GM12878 datasets, we compared the performance (1) using all 440 control datasets against the performance (2) using only 93 control datasets from the related GM cell lines, which have relatively normal karyotypes. Matching control datasets were not used. We observed that the performance slightly increases even when control datasets with abnormal karyotypes were included (Figure S16, p-value=0.02 with Wilcoxon signed rank test). This is expected since our model should be able to put appropriate weights on control datasets, based on how informative they are for imputing common background signal in an IP dataset, especially given that we use 30 million genomic positions as samples which provides strong statistical power. While Figure S12 from the previous section already shows that more control datasets is always better (assuming you don’t know which controls you need), this result on GM cell types shows that even control datasets from abnormal cell types improve the performance on cell types with a normal karyotype thanks to the ability of AIControl to properly integrate background signal structures.

The performance of AIControl indeed dropped when the control datasets from the same cell types are not available. However, it is important to note that our framework, without controls from the same cell type, identified peaks that are better associated with sequence motifs than other peak callers with matched control datasets from the same cell type (Table S5 and S9). This suggests that our framework can successfully estimate the structure of background signals in one cell type by leveraging information from other cell types in a “cross-cell-type” manner.

### AIControl reveals transcription factor interactions better than alternative methods

One of the many downstream use cases of ChIP-seq data is to learn interactions among regulatory factors by observing their co-localization patterns on genome [35, 50, 45]. In particular, Lundberg et al. (2016) [28] showed that the *chromatin network* (i.e., a network of transcription factors (TFs) that co-localize in the genome and interact with each other) can be inferred by estimating the conditional dependence network among multiple ChIP-seq datasets. The authors showed that the inverse correlation matrix computed from a set of ChIP-seq datasets can capture many of the known physical protein-protein interactions (PPIs) from the BioGrid database [42]. Here, we use the same evaluation criteria: significance of the overlap between BioGrid-supported PPIs and the network estimates inferred based on the peaks called by AIControl or by alternative methods. Note that the interactions between datasets that target the same transcription factor and the self-interactions in the diagonal entries are included in this analysis.

Figure 5(a) shows the fold enrichment of true positive predictions over random ones with respect to the number of network edges considered (x-axis) (i.e., sorted based on the magnitude of entries in inverse correlation matrices), as revealed by Lundberg et al. (2016) [28]. Areas under the enrichment curves indicate that AIControl performs better at revealing known PPIs than other methods in K562 (Figure 5). In particular, AIControl ranked more true BioGrid-supported interactions – for example, JUN/STAT1, E2F6/MAX, IRF1/STAT1, and GATA2/JUN – above the threshold (defined as the number of true interactions) than other peak callers (Table S10). Additionally, we performed the same enrichment analysis on the other four cell types: GM12878, HepG2, HeLa-S3, and HUVEC. We observed that the improved performance of AIControl in terms of the area under the enrichment curve consistently generalizes to other cell types (Figure S10). Figure 5(b) visualizes the inverse correlation matrices generated from the peaks called by 5 different peak callers as well as the PPIs documented in BioGrid database (labeled as ‘Truth’). Note that AIControl constructs a chromatin network that best overlaps with the ground truth (i.e., BioGrid PPIs) relative to other methods. Additionally, we compared AIControl against SPP peaks downloaded from ENCODE (“SPP-ENCODE”) and SPP peaks generated with our own pipeline. Similar to the result in the motif enrichment task, AIControl continues to exhibit better performance at recovering PPIs (Figure S17).

**Figure 5.**
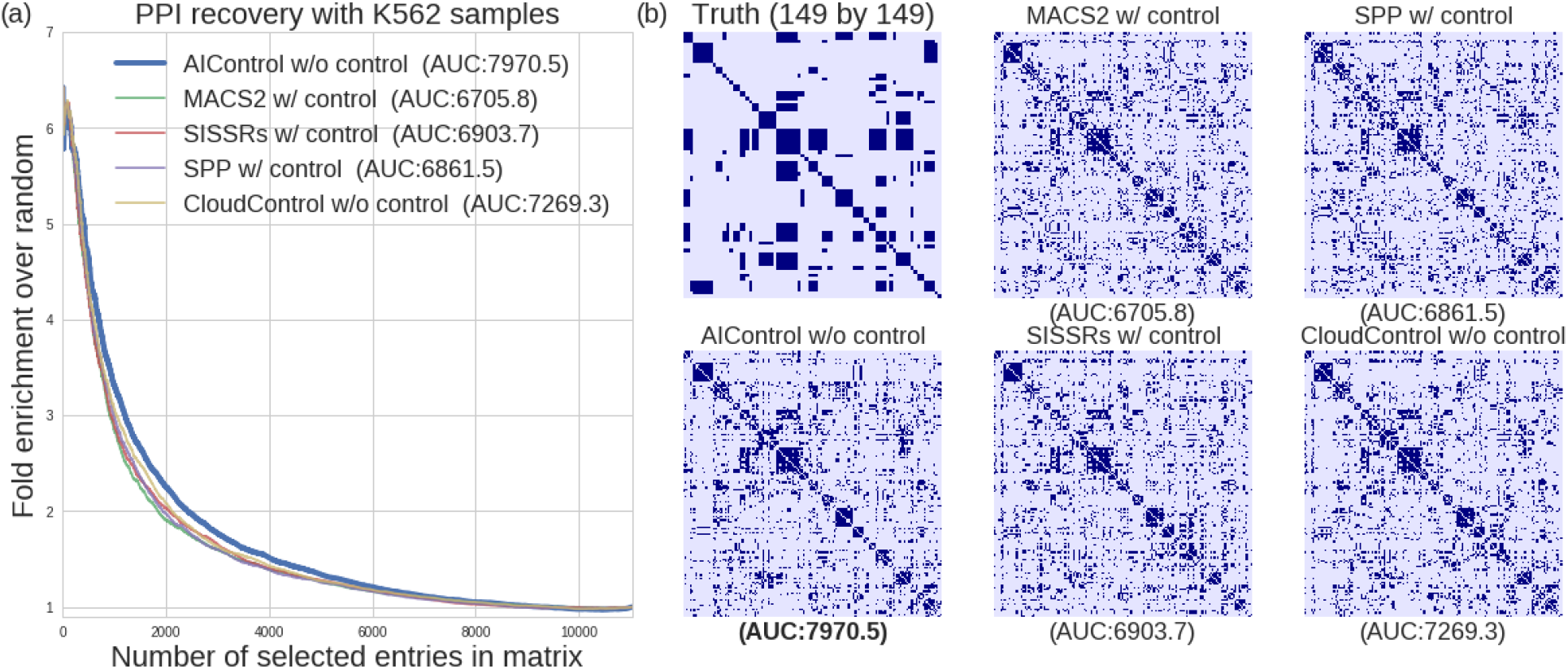
Performance of AIControl compared to other peak callers on the PPI recovery
task in K562. (a) Enrichment of BioGrid-supported interactions of transcription factors in the inverse correlation networks inferred from 149 K562 IP datasets. Peak signals are obtained from: (1) AIControl w/o control, (2) MACS2 w/control, (3) SISSRSs w/control, (4) SPP w/control, and (5) CloudControl + MACS2 w/o control. **(b) Documented interactions among regulatory proteins (top left) and heatmaps of inverse correlation networks (rest)**. The heatmaps are binarized to show top 3583 interactions, which is equal to the number of true interactions.

Table S11 shows top 10 TF interactions that are uniquely suggested by AIControl. Although these interactions are not currently in the database, some studies suggest potential interactions between the pairs. For example, interactions among CEBPB, NFY, and other transcription factors were also thought to play a functionally important roles in the Hypoxia Inducing Factor (HIF) transcriptional response [9]. These predicted interactions, unique to AIControl, may serve as potential targets for discovering previously uncharacterized PPIs.

Although we showed that AIControl better recovers known PPIs, it is important to note that the truth matrix likely contains some false positives and potentially many false negatives. First, the truth matrix is constructed using information drawn from all available cell types. Second, some interactions might still be undocumented in the BioGrid database. Further, our prediction from ChIP-seq data is more likely to recover interactions near DNA strands. Despite these uncertainties, the finding that AIControl recovers PPIs more accurately in all cell types suggests that using this framework can improve the quality of downstream analysis that follows ChIP-seq experiments.

### AIControl better removes common background signal among datasets

One of the most frequently used quality measures for biological experiments is the *consistency* of a pair of replicate datasets, which can be measured by the number of shared peaks. A pair of replicate datasets should capture the exact same signals; thus, a pair with better quality should share more peaks with each other. On the other hand, the quality of background signal removal can be assessed by measuring the *inconsistency* in a pair of unrelated datasets. We define an “unrelated” pair as a pair of datasets that (1) is in the same cell type and (2) targets unrelated transcription factors without any documented PPI in BioGrid. As described in Methods, AIControl models ChIP-seq experiments as follows:

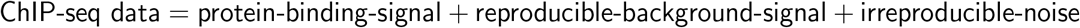

For a pair of unrelated datasets, we assume that there is no `protein-binding-signal` that give rise to shared peaks. The only source of shared peaks in an unrelated pairs is `reproducible-background-signal. If` a peak caller perfectly removes `reproducible-background-signal`, it should ideally leave no peak that is shared between a pair of unrelated datasets. Thus, peak callers better able to remove common background signal should have fewer shared peaks for unrelated datasets. This metric alone is not perfect, because a pair of completely random peaks can achieve good results as well. However, we believe that, in combination with the motif enrichment and PPI recovery task, this metric highlights an important aspect of noise removal process.

Figure S18 shows the “sharedness” of peaks for 9,310 pairs of unrelated datasets in K562 processed by five peak callers: AIControl, MACS2, SISSRs, SPP, and CloudControl+MACS2. The y-axis indicates the proportion of unrelated datasets that have less than a particular number of shared peaks, which is represented in the x-axis. The smaller area under the curve demonstrates that a peak caller generally identifies fewer shared peaks between a pair of unrelated datasets, and AIControl exhibits the smallest area under its curve. For other peak callers, a larger percentage of unrelated dataset pairs contained more shared peaks, suggesting that their bias-removal process was not as thorough as that of AIControl’s.

### Exploring the compatibility of AIControl and other peak callers with the IDR method

We investigated the performance of AIControl in a situation where biologically replicated samples are available. In particular, the ENCODE consortium uses the irreproducible discovery rate (IDR) framework to adaptively rank and select peaks based on the rank consistency/reproducibility of signals among biological replicates [26]. The ENCODE official pipeline uses SPP in combination with IDR to identify and reorder peaks in ChIP-seq datasets. In order to evaluate the effect of IDR on the peak callers, for datasets where biological replicates are available, we performed IDR analysis after calling peaks with AIControl, MACS2, and SISSRs. For SPP, we directly downloaded peaks processed with IDR from the ENCODE website.

Figure S1 (a) shows the superior performance of AIControl+IDR in the motif sequence identification task across five tested cell types (i.e., K562, GM12878, HepG2, HeLa-S3, and HUVEC) compared to that of other peak callers with IDR. Negative log10 idr values were used as a ranking measure. Needless to say, other peak callers have access to matched control datasets, whereas AIControl does not. We also observed that AIControl+IDR better predicts protein-protein interactions in the K562 cell type than other peak callers when they are used in combination with IDR (Figure S1 (b)).

### AIControl performs well on the datasets outside the ENCODE database

The recommended protocol strictly regulates ChIP-seq experiments in the ENCODE database. However, external labs do not always adhere precisely to this protocol. To assure that the AIControl framework generalize strong performance on a ChIP-seq IP dataset that is not a part of the ENCODE database, we performed peak calling on 14 IP datasets that are obtained from 8 independent studies [27, 11, 38, 40, 49, 29, 46, 37], which are not part of the ENCODE database and are only on the GEO or ArrayExpress database. We only analyzed the datasets whose target motif PWMs for *H. sapiens* are available in the JASPAR database. The AUPRC values were measured across the whole genome. The information of the 14 datasets are summarized in Table S4. We compared the following peak calling frameworks: AIControl, MACS2, SISSRs, and SPP.

Figure 6 shows the performance of AIControl on all 14 external datasets in comparison to other peak callers. Individual PR curves are shown in Figure S19. AIControl retains its strong performance on all datasets except for the one that targets SP1 in the HEK293 cell type. This is consistent with the result shown in Figure 2 (a), where AIControl exhibits the worst performance on the datasets with SP1.

**Figure 6.**
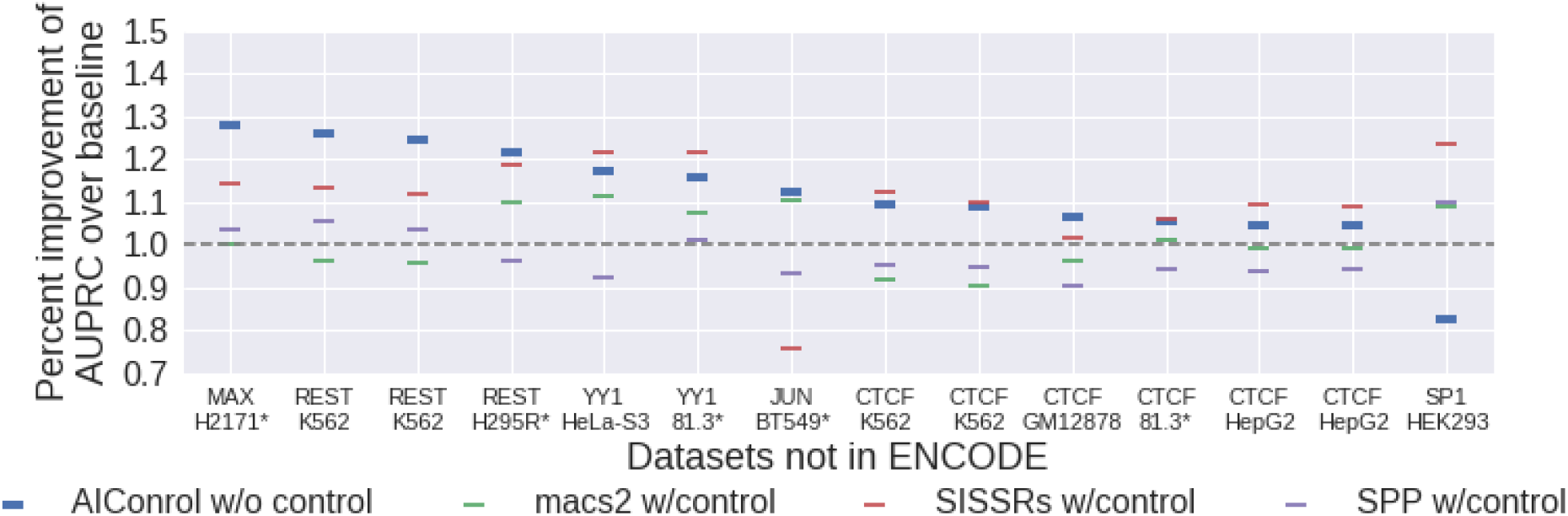
Relative performance of five peak calling methods on external datasets. Peak regions were identified using: (1) AIControl w/o control, (2) MACS2 w/control, (3) SISSRSs w/control, and (4) SPP w/control. The y-axis shows the percent improvement of the area under the precision-recall curves (AUPRCs) for predicting the presence of putative binding sites with significance values associated with the peaks over the baseline (i.e., MACS2 without using a matched control dataset). The x-axis shows the non-ENCODE ChIP IP datasets ordered by the percent improvement achieved by AIControl (y-axis). Datasets measured in celltypes that are not in the 440 ENCODE control set are shown with asterisks.

Most notably, 5 out of the 14 external datasets were measured in the cell types that were not included in the pool of control datasets used for AIControl. All datasets had matched control datasets, which were used by MACS2, SISSRs, and SPP, but not by AIControl. The fact that AIControl without matched control datasets retained strong performance suggests that it is able to integrate background control signals in a cross-cell-type setting even in a case where the datasets are not as highly regulated as that of ENCODE database.

### Principal components are associated with potential bias sources

Control datasets are supposed to capture background signals that are also present in corresponding IP datasets. Many studies suggest that these background signals are combinations of multiple different sources of biases, for instance, GC content, sonication bias, and platform-specific biases [36, 21, 25]. The AIControl framework assumes that observed background signals in a control dataset can be represented as the weighted sum of many different known or unknown bias sources (see Methods).

Figure S20 shows Spearman’s correlation coefficients between potential bias sources and K562 control datasets projected on the first five principal components. Open chromatin regions (HS) and read mappability (MP) are similar to the first principal component, while GC content (GC) is similar to the second principal component. Notably, the first five principal components collectively capture only 54.5% variance, which suggests other bias sources are likely to exist that contribute to the observed background signal. AIControl implicitly learns the contributions from unobserved sources of biases; this is one of the reasons that AIControl can call more accurate peaks relative to other peak identification methods.

## Discussion

Accurately identifying the locations of regulatory factor binding events remains a core, unresolved problem in molecular biology. AIControl offers a framework for processing ChIP-seq data to identify binding locations of transcription factors without requiring a matched control dataset.

AIControl makes key innovations over existing systems. (1) It learns position-specific distribution of background signal at much finer resolution than other methods by using publicly available control datasets on a large scale (see Methods). Our evaluation metrics show that using finer background distributions improved enrichment of putative TF binding locations and recovery of known protein-protein interactions. (2) AIControl systematically integrates control datasets from a public database (e.g., ENCODE) without any user input. Its ability to learn background signals extends to datasets obtained in new cell types without any previously measured control datasets. We obtained 440 ChIP control datasets from 107 cell types in the ENCODE database, and AIControl learns to statistically combine them to estimate background signals in an IP dataset in any cell type. We showed that our performance on new cell types exceeds that of established baselines. AIControl’s performance is also generalizable to datasets from labs outside the ENCODE project. (3) The mathematical model of AIControl accounts for multiple sources of biases due to its integration of control datasets at a large scale (see Methods). On the other hand, some sources of biases may not be fully captured by existing methods that use only one matched control dataset [48, 33] or account for only a specific set of biases. [21, 36]. (4) Finally, AIControl reduces the time and cost incurred by generating a matched control dataset since it does not require a control to perform rigorous peak calling.

We demonstrated the effectiveness of AIControl by conducting a large-scale analysis on the peaks identified in the 410 ENCODE ChIP-seq datasets from five major ENCODE cell types for 54 different transcription factors (Table S1). We showed that AIControl has better motif sequence enrichment compared to other peak callers within predicted peak locations. However, this metric measures only direct interactions between transcription factors and DNA. Thus, we evaluated the performance of AIControl with another metric: PPI enrichment analysis. In this metric, we also observed that AIControl is superior to other peak callers even without any matched control samples. In conclusion, we showed that our framework’s single-dataset peak identification performs better than other established baselines with matched controls datasets.

AIControl satisfies many of the properties favored by the comparative analysis of peak calling algorithms [43, 47, 23]. This includes the use of local distributions that are suitable for modeling count data and the ability to combine ChIP-seq and input signals in a statistically principled manner. There are several future extensions for our framework. (1) Our default implementation bins IP and control datasets into 100 bp windows in order to perform fast genome-wide regression. Because most transcription factors show signals wider than 100 bps, we believe that our resolution is sufficient to conduct accurate downstream analysis. The Julia implementation can be accessed at https://github.com/hiranumn/AIControl, and the accompanying files can be accessed through the Google Drive link under the “Paper” section on the GitHub page. (2) Since our framework learns weights that are globally applied to all genomic positions, it performed relatively worse on estimating background signal for cell types with abnormal karyotype in a cross-cell-type setting. This could be resolved by automatically detecting karyotype abnormality and learning different sets of weights for those regions. (3) We currently provide control signals that are mapped to the hg38 human genome assembly on our website. Therefore, our framework also expects ChIP-seq datasets in the hg38 space. If an user wants to use our framework for identifying peaks in the hg19 genome assembly, one can currently use the UCSC liftover tool to convert the called peaks from the hg38 to hg19 space.

ChIP-seq is one of the most widely used techniques for identifying protein binding locations. However, conducting a set of two ChIP-seq experiments can be resource intensive. By removing the cost of obtaining control datasets, we believe that AIControl can lead to more accurate ChIP-seq signals without expending additional resources.

## Supplementary Data

Supplementary Data are available online.

### Funding

National Science Foundation (DBI-1552309).

### Supplementary Information

**Figure S1.**
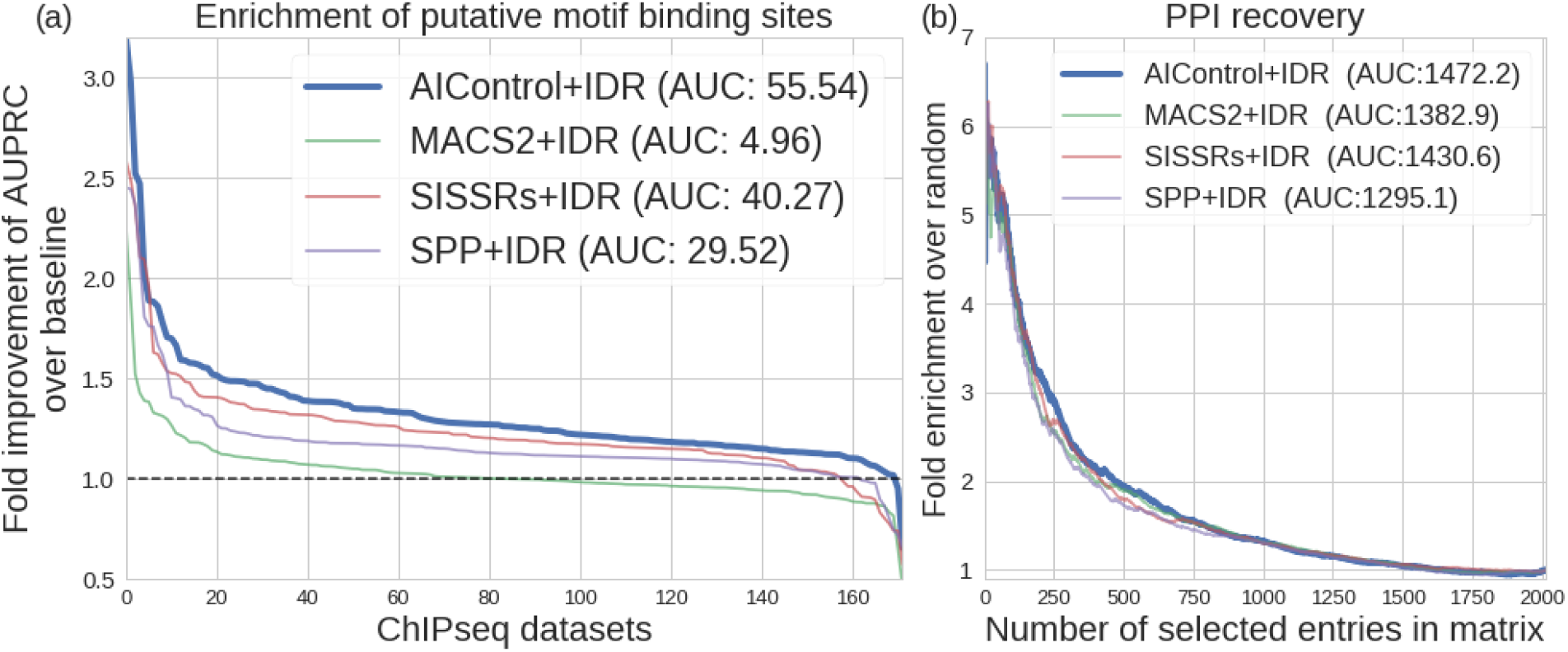
(a) Relative performance of peak calling methods compared to MACS2 without using a matched control dataset as a baseline (dotted line) in combination with IDR. Peaks were identified using: (1) AIControl+IDR w/o control, (2) MACS2+IDR w/control, (3) SISSRSs+IDR w/control, and (4) SPP+IDR w/control. The y-axis shows the fold improvement of the area under the precision-recall curves (AUPRCs) for predicting the presence of putative binding sites with significance values associated with the peaks over the baseline (i.e., MACS2 without using a matched control dataset) across the whole genome. The x-axis shows the all 168 biological replicate pairs of ENCODE ChIP IP datasets ordered by the fold improvement (y-axis). Note that the ordering of datasets is different for each peak caller. Area between each line and the dotted baseline is shown in parenthesis. **(b) Performance of AIControl+IDR compared to other peak callers on the PPI recovery task in K562**. Enrichment of BioGrid-supported interactions of transcription factors in the inverse correlation networks inferred from 64 K562 IP datasets.

**Figure S2.**
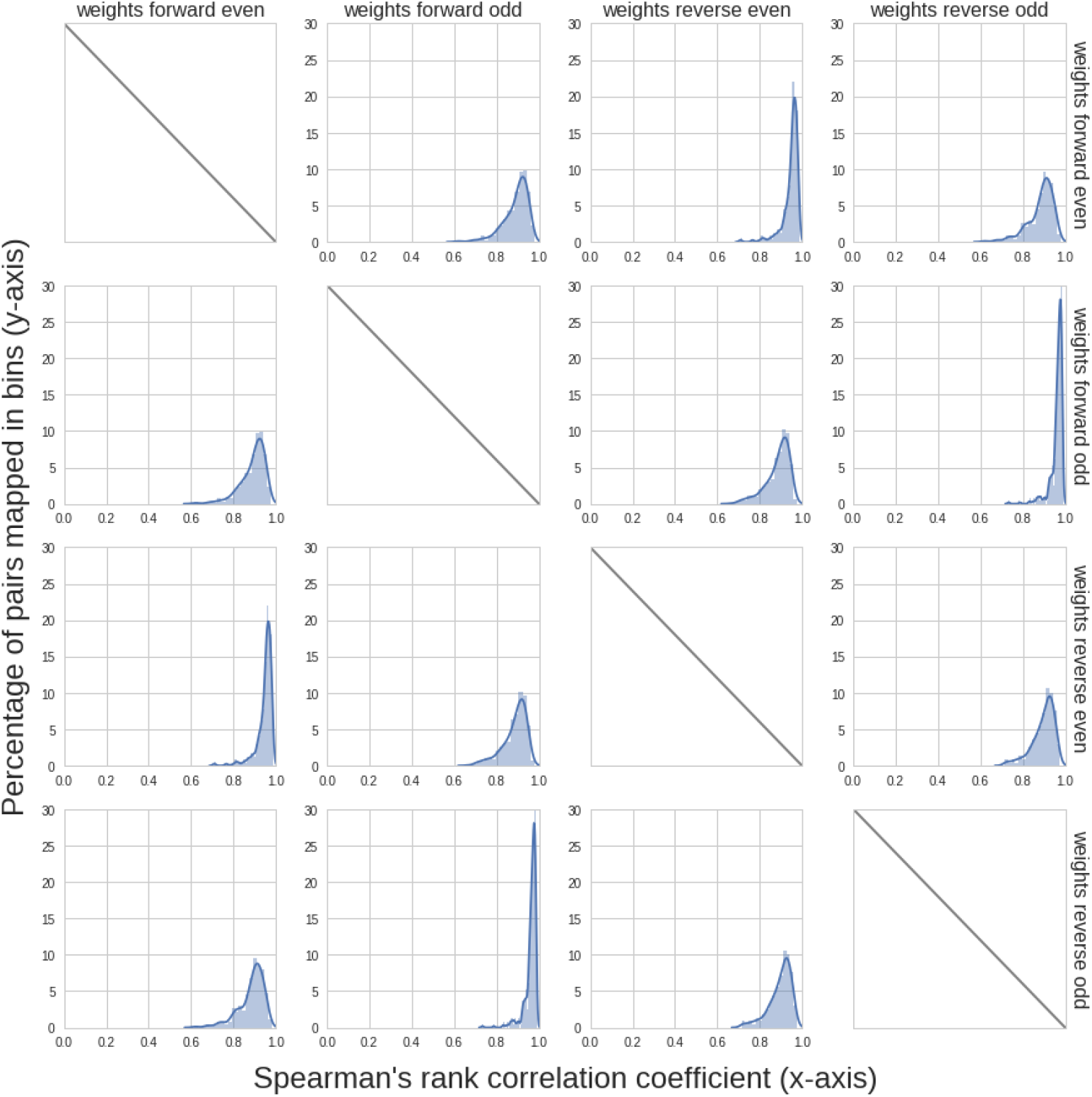
A histogram of Spearman correlation values for all possible pairs among 4 weight vectors learned across all IP datasets. The smooth lines are generated by performing kernel density estimates with bandwidth determined by Scott’s rule [41].

**Figure S3.**
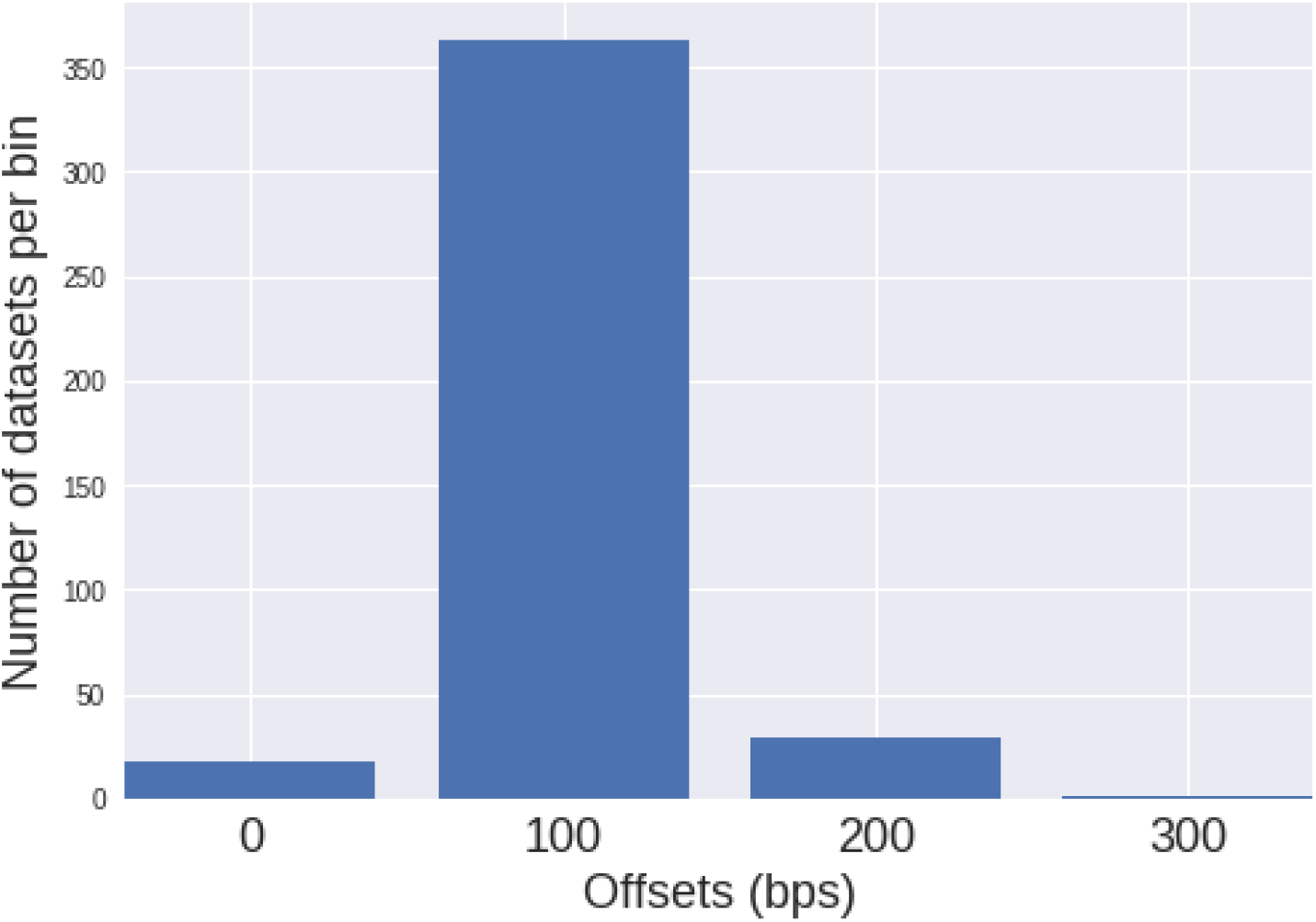
A histogram of automatically estimated distance between reverse and forward peaks (i.e., *d*). Most datasets have *d* = 100, which is expected for normal ChIP-seq experiments for transcription factors.

**Figure S4.**
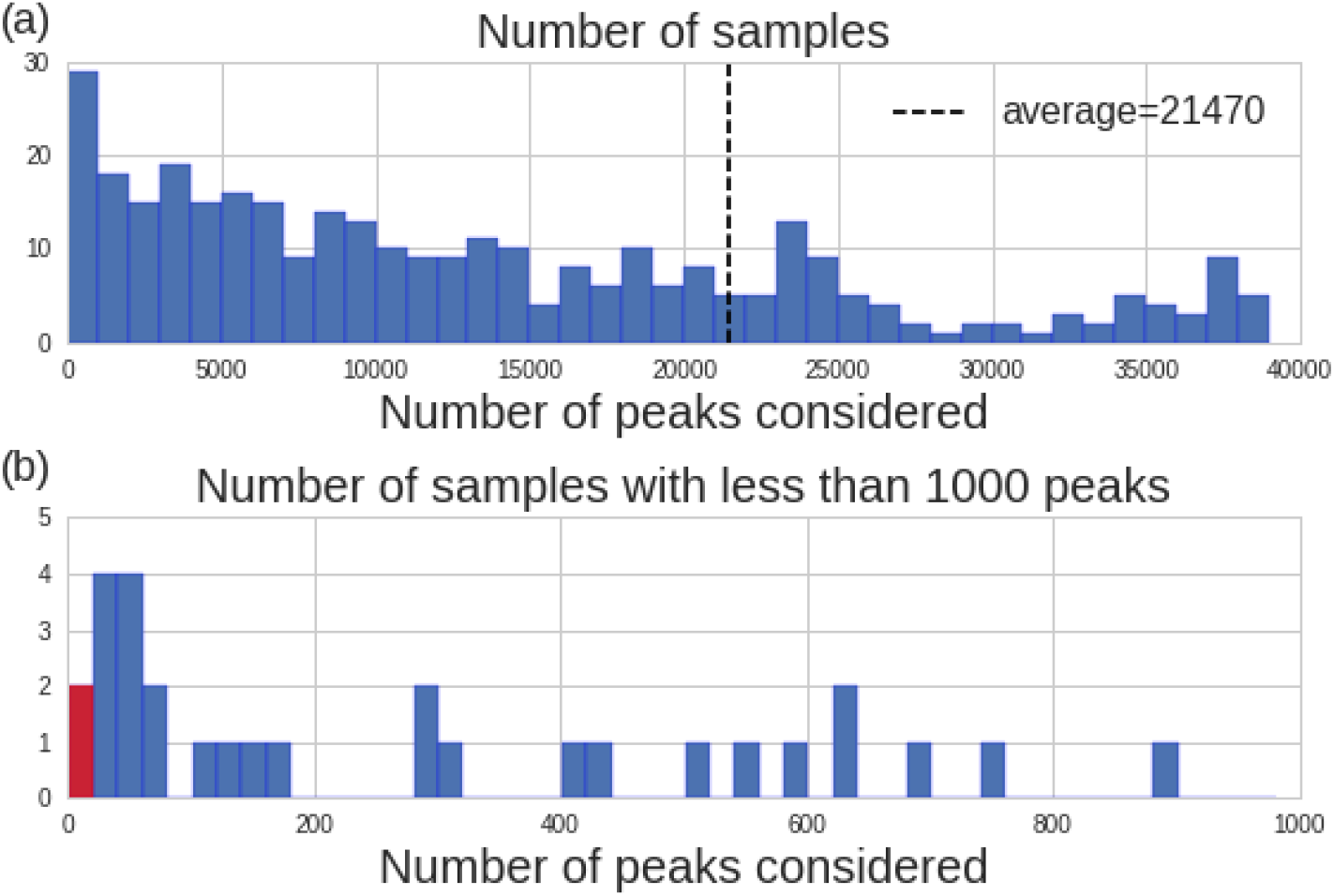
(a) A histogram of the numbers of peaks considered by our evaluations across the whole genome. For each IP dataset, the minimum number of peaks identified among the five peak callers was used for downstream evaluations (see Methods, “Standardizing peak signals”) **(b) A histogram showing the distribution of the number of peaks in the first bar of** (a) (corresponds to 0~1000 peaks). The red bar indicates the datasets omitted in Figure 2 for the better stability of area under PR curves.

**Figure S5.**
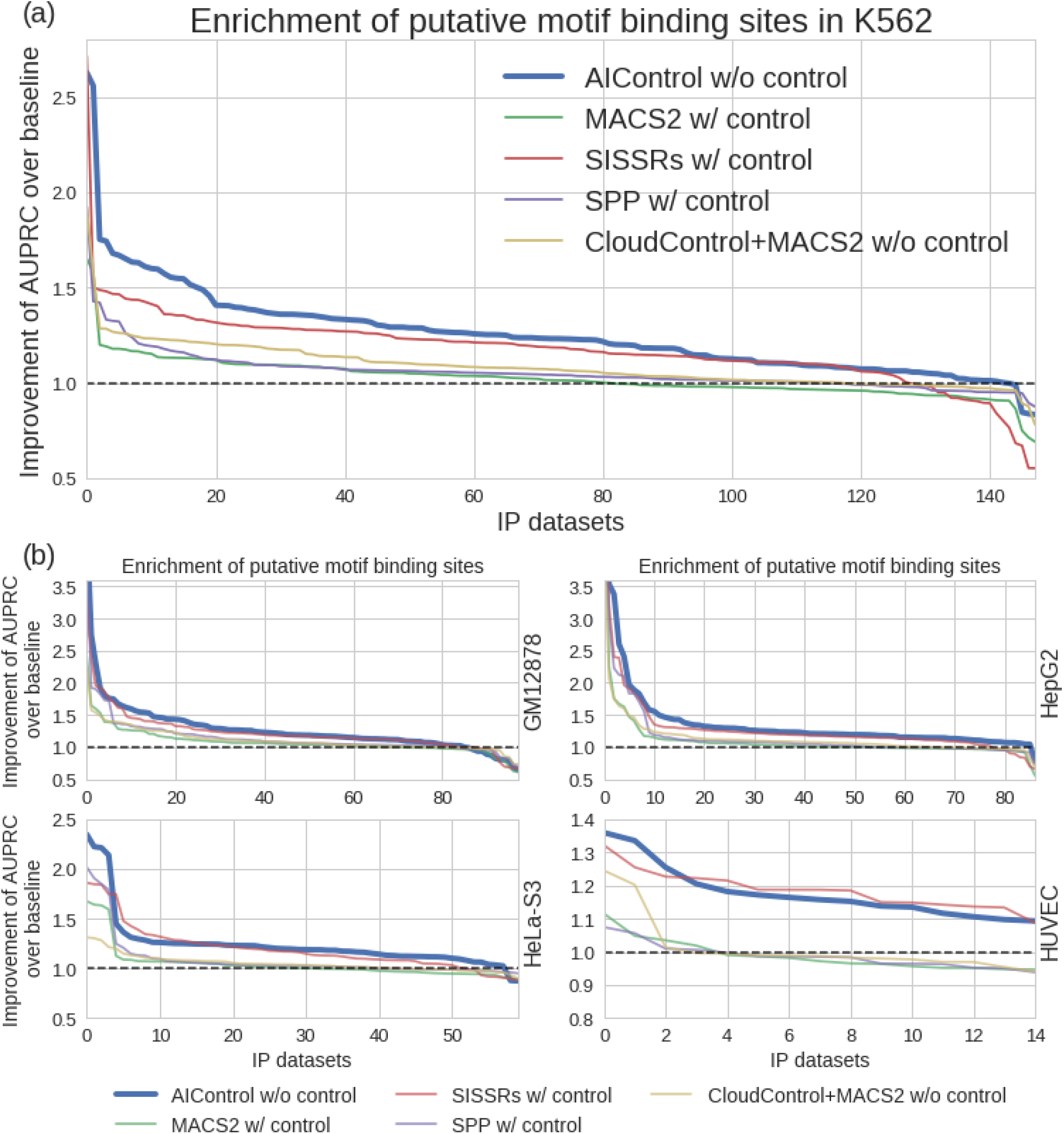
Relative performance of five peak calling methods compared to MACS2 without using a matched control dataset as a baseline (dotted line) across the whole genome. Peaks were identified using: (1) AIControl w/o control, (2) MACS2 w/control, (3) SISSRSs w/control, (4) SPP w/control, and (5) CloudControl + MACS2 w/o control. The y-axis shows the fold improvement of the area under the precision-recall curves (AUPRCs) for predicting the presence of putative binding sites with significance values associated with the peaks over the baseline (i.e., MACS2 without using a matched control dataset) across the whole genome. The x-axis shows the ENCODE ChIP IP datasets in each cell type ordered by the fold improvement (y-axis) for: (a) 149 IP datasets from K562, and (b) other tier 1 ENCODE cell types (i.e., GM12878, HepG2, HeLa-S3, and HUVEC). Note that the ordering of datasets is different for each peak caller.

**Figure S6.**
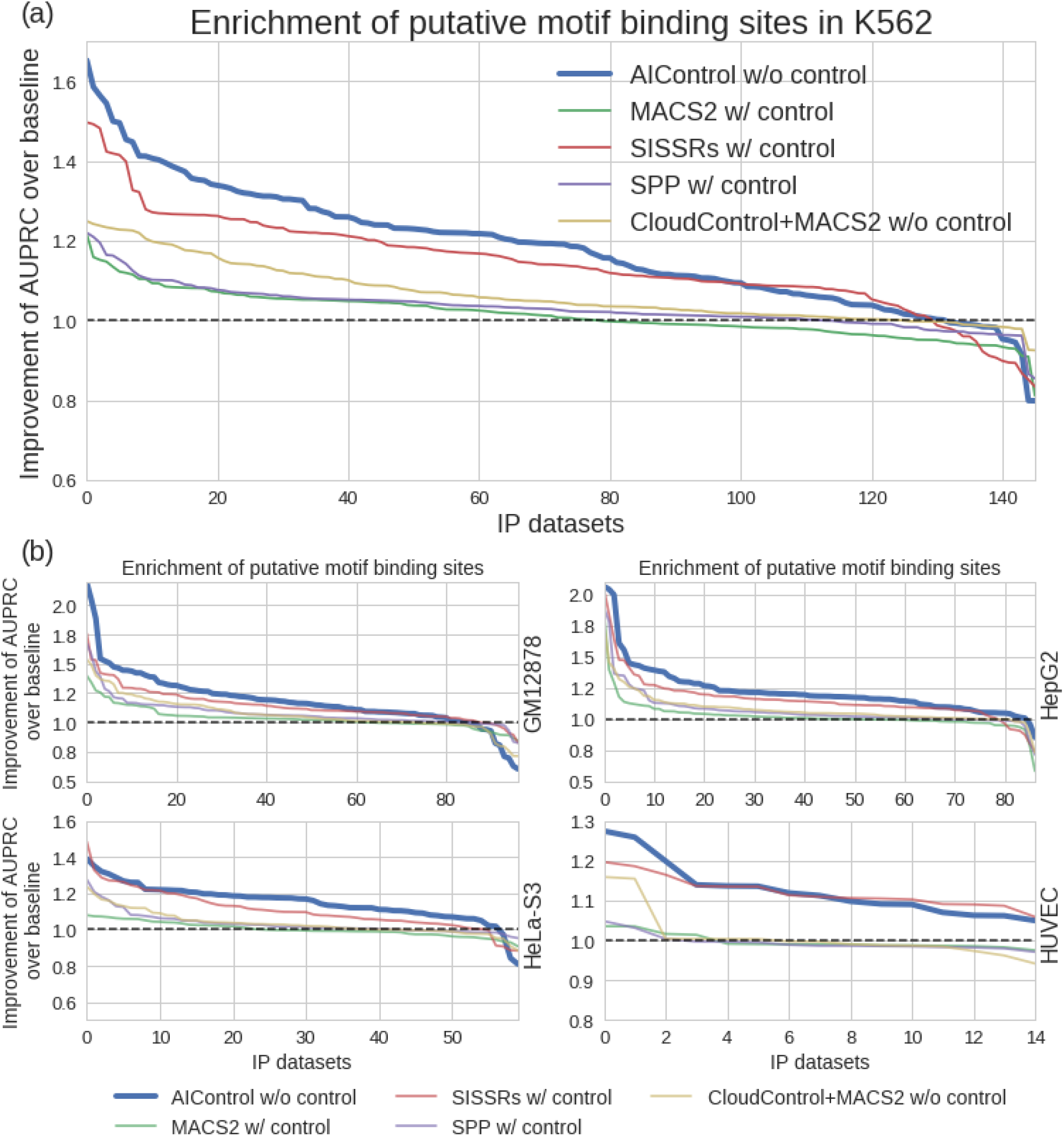
Relative performance of five peak calling methods compared to MACS2 without using a matched control dataset as a baseline (dotted line) in the DNase 1 hypersensitivity regions (DHS). Peaks were identified using: (1) AIControl w/o control, (2) MACS2 w/control, (3) SISSRSs w/control, (4) SPP w/control, and (5) CloudControl + MACS2 w/o control. The y-axis shows the fold improvement of the area under the precision-recall curves (AUPRCs) for predicting the presence of putative binding sites with significance values associated with the peaks over the baseline (i.e., MACS2 without using a matched control dataset) in the DHS. The x-axis shows the ENCODE ChIP IP datasets in each cell type ordered by the fold improvement (y-axis) for: (a) 146 IP datasets from K562, and (b) other tier 1 ENCODE cell types (i.e., GM12878, HepG2, HeLa-S3, and HUVEC). Note that the ordering of datasets is different for each peak caller.

**Figure S7.**
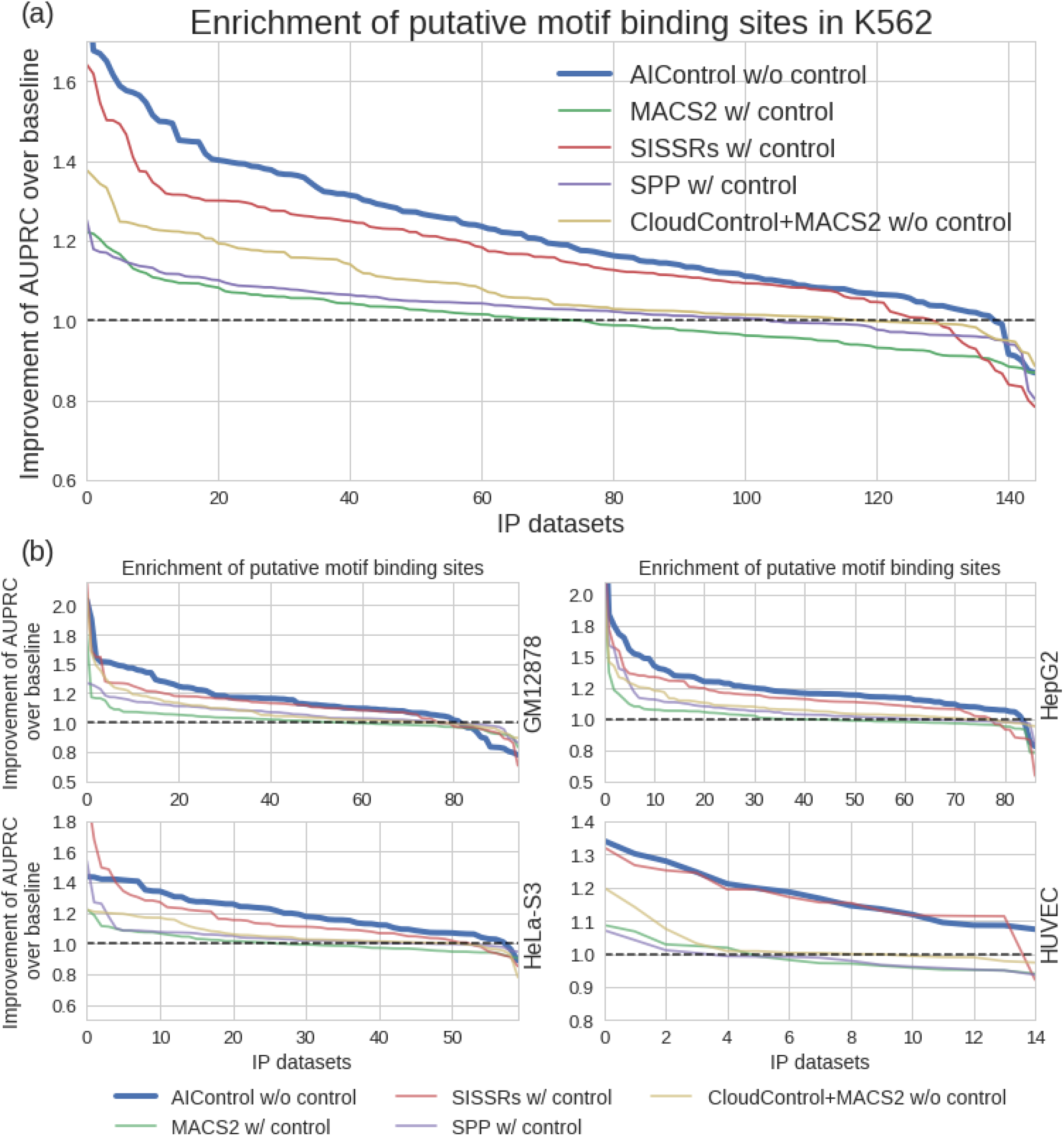
Relative performance of five peak calling methods compared to MACS2 without using a matched control dataset as a baseline (dotted line) in regions 5000 bps up and downstream of protein coding gene start sites. Peaks were identified using: (1) AIControl w/o control, (2) MACS2 w/control, (3) SISSRSs w/control, (4) SPP w/control, and (5) CloudControl + MACS2 w/o control. The y-axis shows the fold improvement of the area under the precision-recall curves (AUPRCs) for predicting the presence of putative binding sites with significance values associated with the peaks over the baseline (i.e., MACS2 without using a matched control dataset) near the protein coding regions. The x-axis shows the ENCODE ChIP IP datasets in each cell type ordered by the fold improvement (y-axis) for: (a) 145 IP datasets from K562, and (b) other tier 1 ENCODE cell types (i.e., GM12878, HepG2, HeLa-S3, and HUVEC). Note that the ordering of datasets is different for each peak caller.

**Figure S8.**
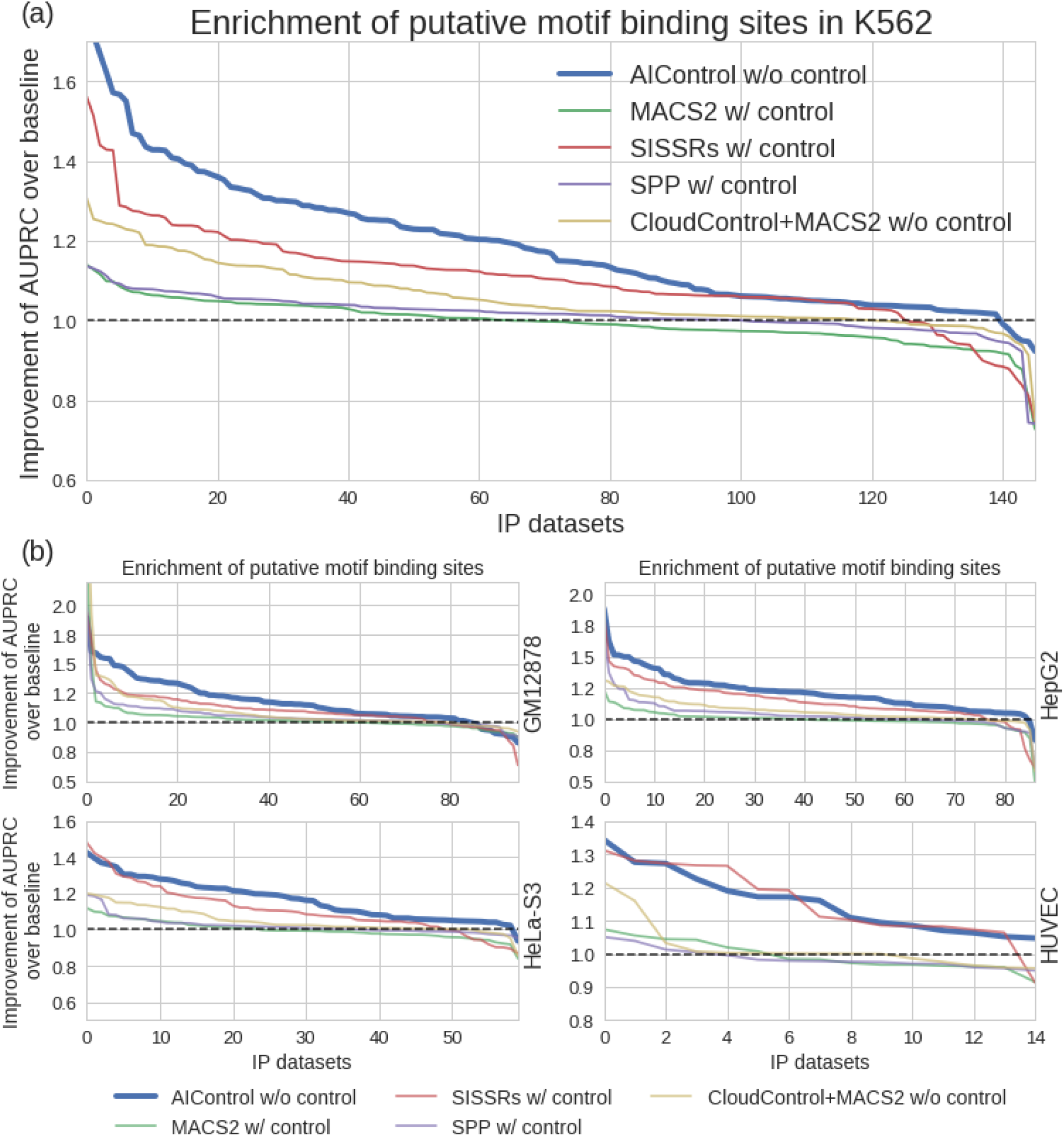
Relative performance of five peak calling methods compared to MACS2 without using a matched control dataset as a baseline (dotted line) in regions with more than 50% GC content. Peaks were identified using: (1) AIControl w/o control, (2) MACS2 w/control, (3) SISSRSs w/control, (4) SPP w/control, and (5) CloudControl + MACS2 w/o control. The y-axis shows the fold improvement of the area under the precision-recall curves (AUPRCs) for predicting the presence of putative binding sites with significance values associated with the peaks over the baseline (i.e., MACS2 without using a matched control dataset) in the GC rich regions. The x-axis shows the ENCODE ChIP IP datasets in each cell type ordered by the fold improvement (y-axis) for: (a) 145 IP datasets from K562, and (b) other tier 1 ENCODE cell types (i.e., GM12878, HepG2, HeLa-S3, and HUVEC). Note that the ordering of datasets is different for each peak caller.

**Figure S9.**
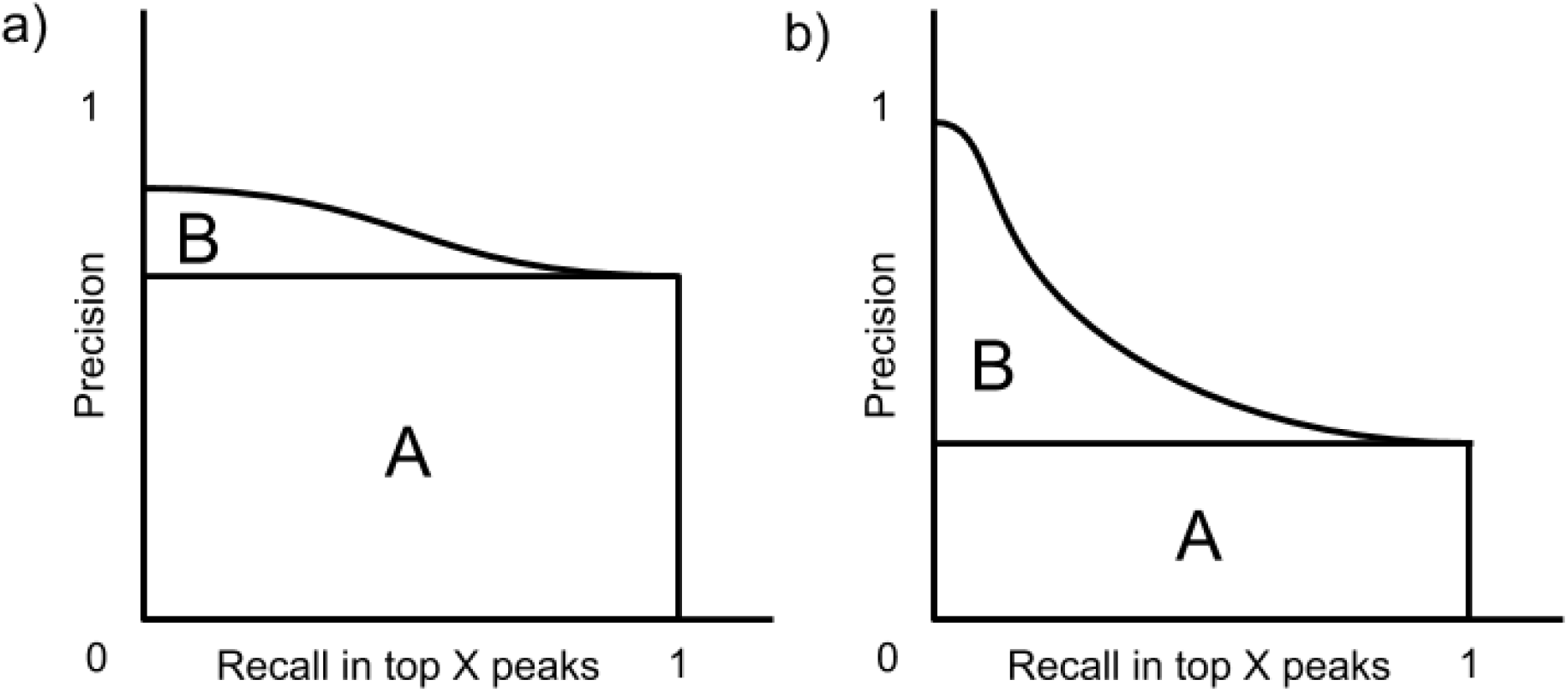
(a) An example PR curve for a peak caller that performs well at selecting true peaks in top *n* (captured by area A) but poorly at ordering them in top *n* (captured by area B). (b) An contrasting example PR curve for a peak caller that performs poorly at placing true peaks in top *n* but well at ordering them in top *n*. Both area A and B are important for high quality peak calling.

**Figure S10.**
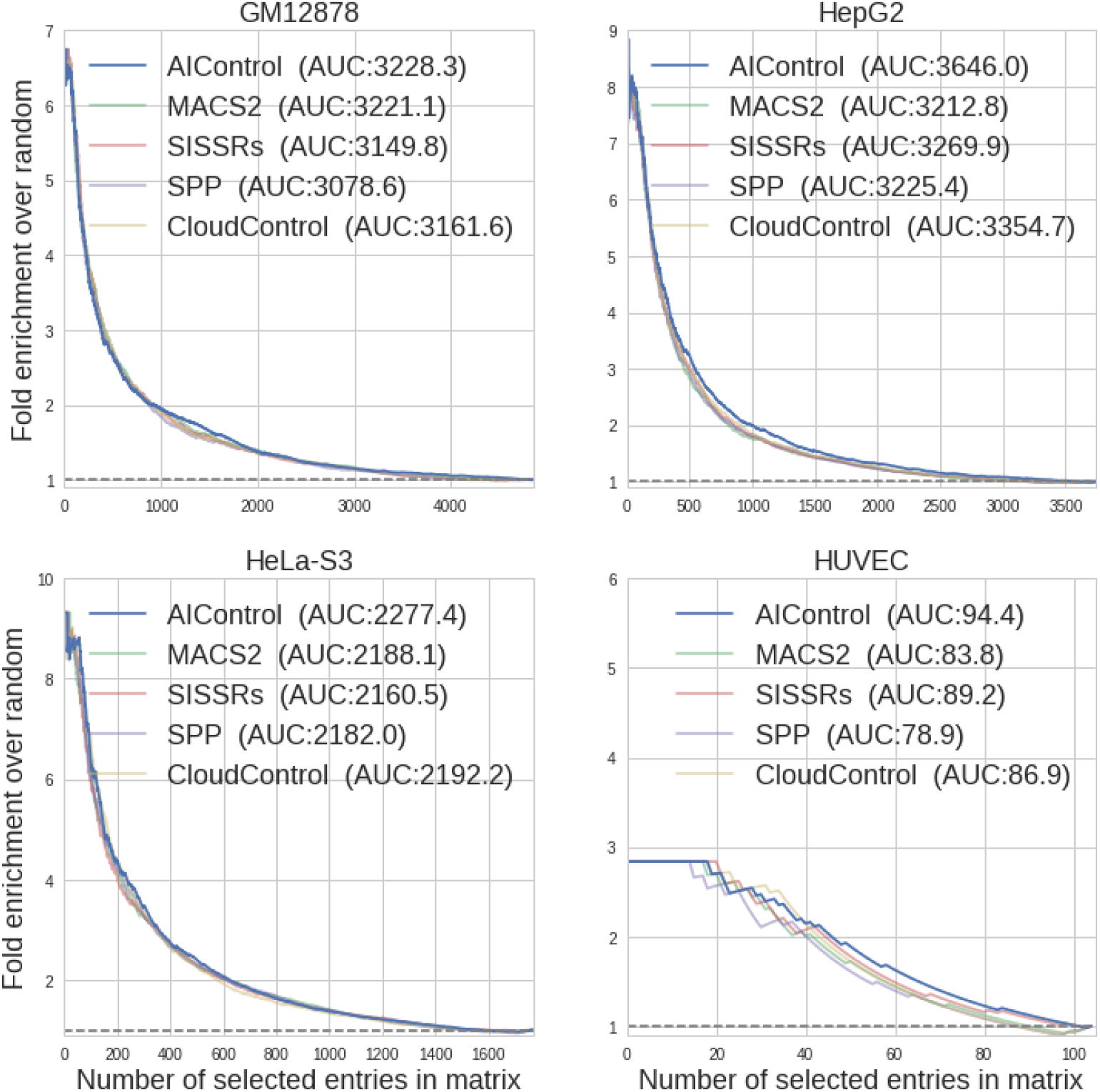
Performance of AIControl compared to other peak callers on the PPI recovery task in the cell types other than K562. Each plot shows enrichment of BioGrid-supported interactions between transcription factors in the inverse-correlation networks inferred from peak signals obtained from five different peak calling frameworks (i.e., AIControl w/o control, MACS2 w/control, SISSRSs w/control, SPP w/control, and CloudControl + MACS2 w/o control) in four ENCODE tier 1 or 2 cell lines (i.e., GM12878, HeLa-S3, HepG2, and HUVEC).

**Figure S11.**
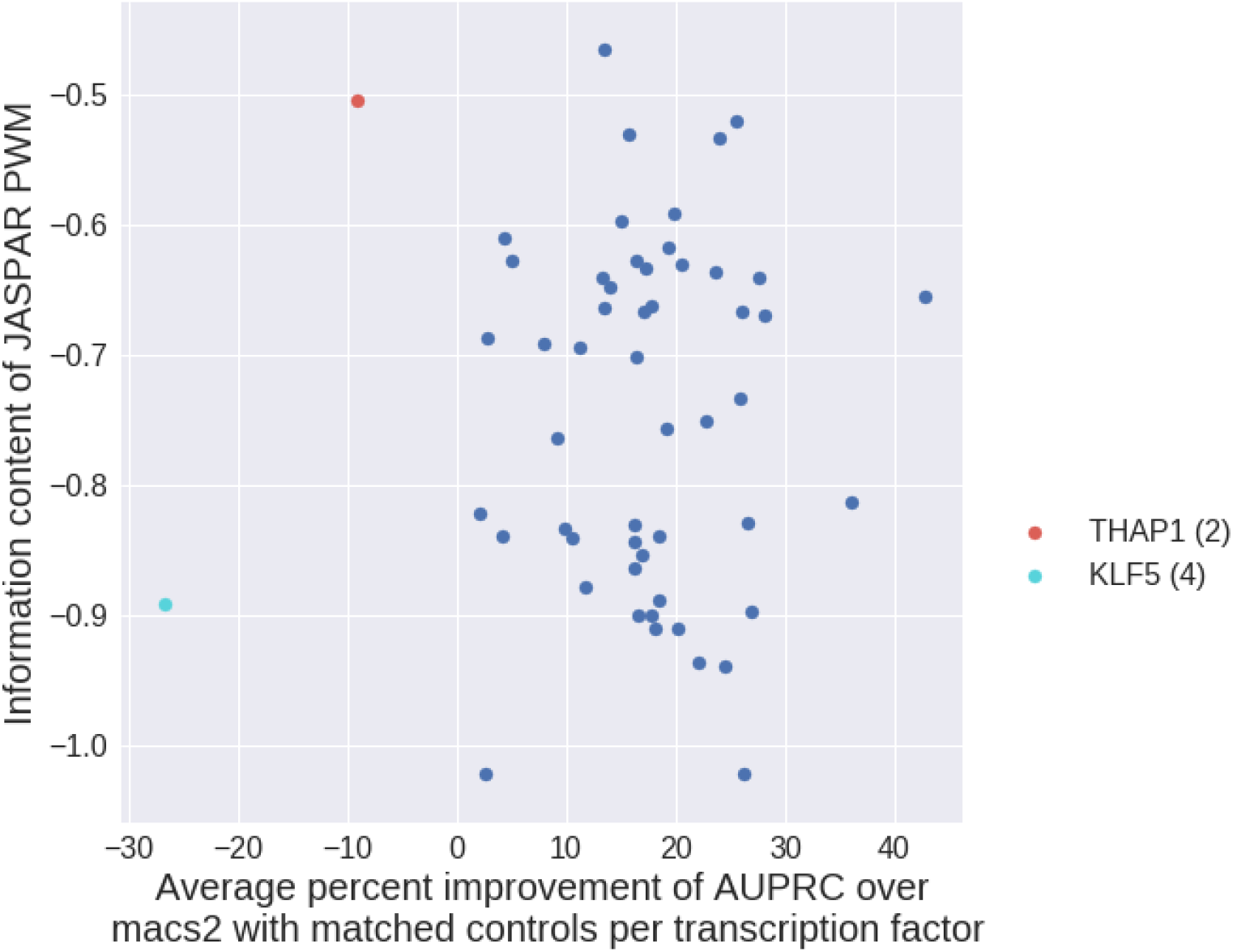
Information content (sum of mean relative entropy per base pair position) of PWMs plotted against the ratio of improvement from MACS2 to AIControl on the AUPRC metric for predicting transcription factor binding events (r=0.02, p-value=0.84). Each point represents a transcription factor. In particular, AIControl performed worse on the following regulatory factors: THAP1, KLF5. The numbers inside parentheses indicates the number of datasets that target corresponding factors.

**Figure S12.**
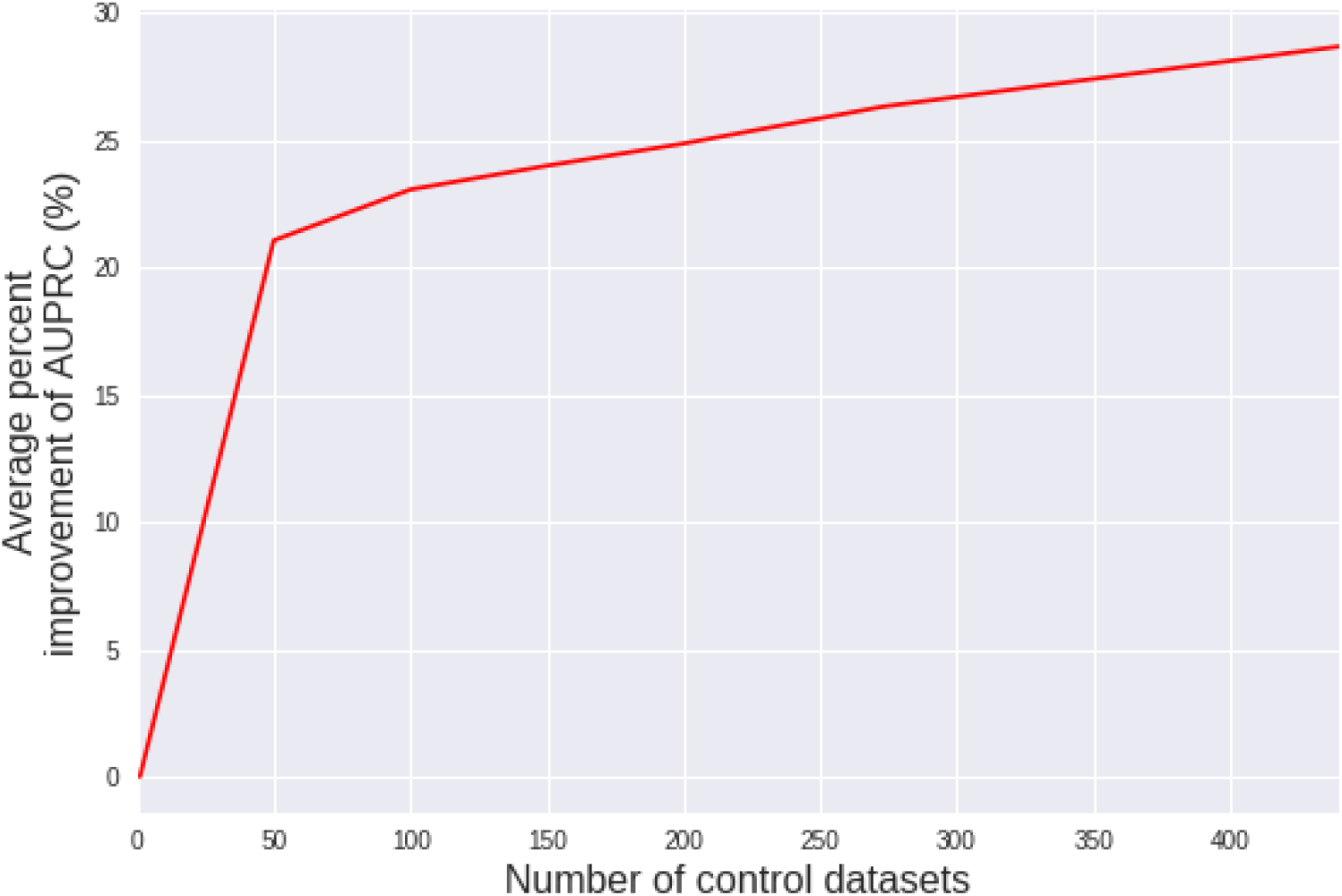
Average percent change in AUPRCs with respect the number of controls used by our AIControl framework. The plot shows the average behavior of 36 ChIP-seq IP experiments from 5 cell types (K562, GM12878, HepG2, HeLa-S3, HUVEC) for TFs that are measured more than 10 times. Some TF/cell-line pairs are missing due to data availability. For each IP experiment at each subset size, the AUPRC values were averaged over the number of random subsets equal to ceil(450/(subset size)). For example, to measure the performance with a subset of 50 control datasets, we selected random subsets 9 times.

**Figure S13.**
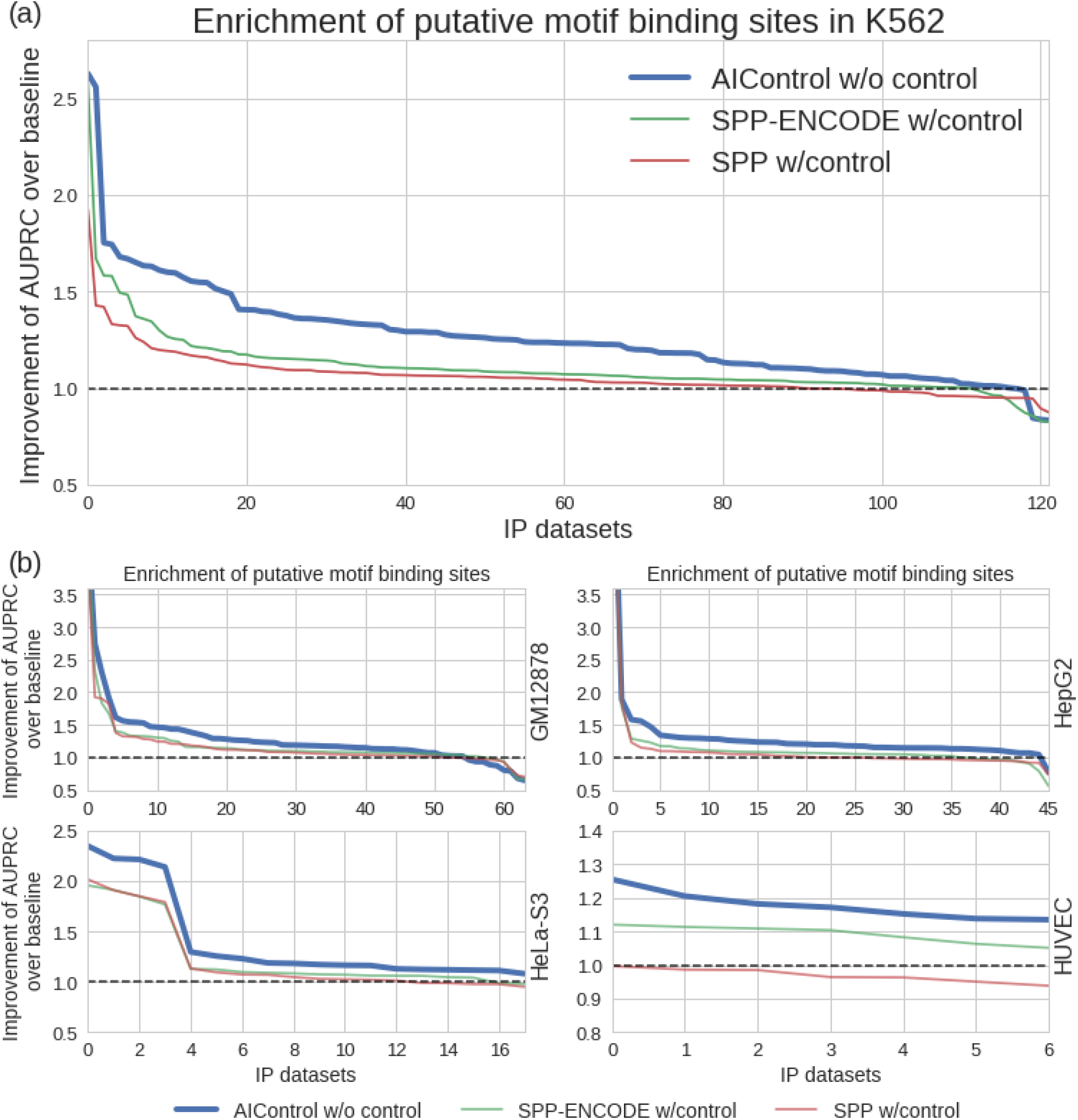
Relative performance of three peak calling methods compared to MACS2 without using a matched control dataset as a baseline (dotted line) across the whole genome. Peaks were identified using: (1) AIControl w/o control, (2) SPP peaks from the ENCODE database (“SPP-ENCODE”), and (3) SPP w/control. The y-axis shows the fold improvement of the area under the precision-recall curves (AUPRCs) for predicting the presence of putative binding sites with significance values associated with the peaks over the baseline (i.e., MACS2 without using a matched control dataset). The x-axis shows the ENCODE ChIP IP datasets in each cell type ordered by the fold improvement (y-axis) for: (a) 122 IP datasets from K562, and (b) other tier 1 ENCODE cell types (i.e., GM12878, HepG2, HeLa-S3, and HUVEC). Note that the ordering of datasets is different for each peak caller.

**Figure S14.**
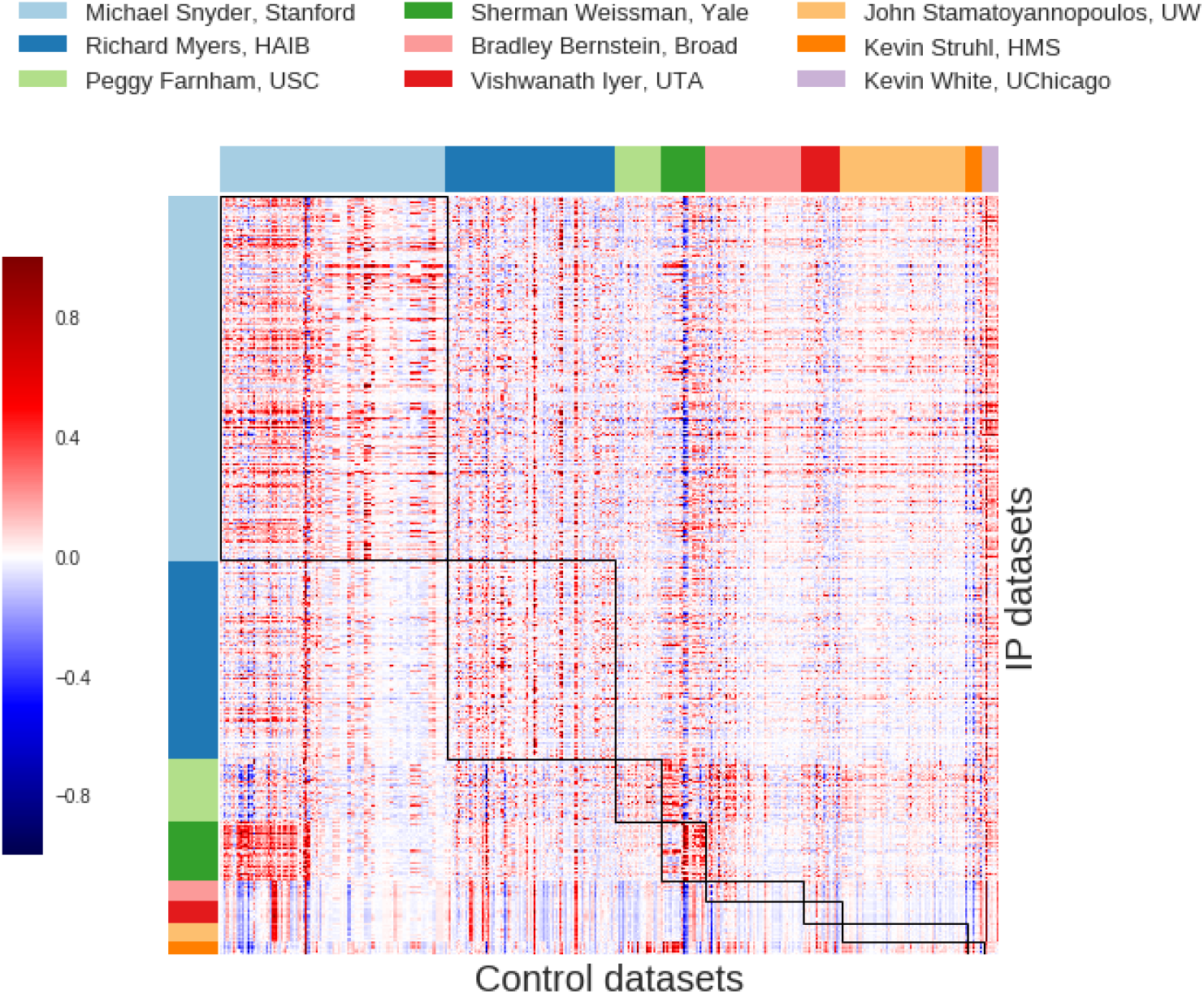
Normalized weights the AIControl model assigns to 440 ENCODE control datasets (columns) for each of the 410 IP datasets (rows) sorted by labs. (Tables S1 and S2). The black rectangles indicate the weights for the control datasets measured in the same lab as the IP datasets.

**Figure S15.**
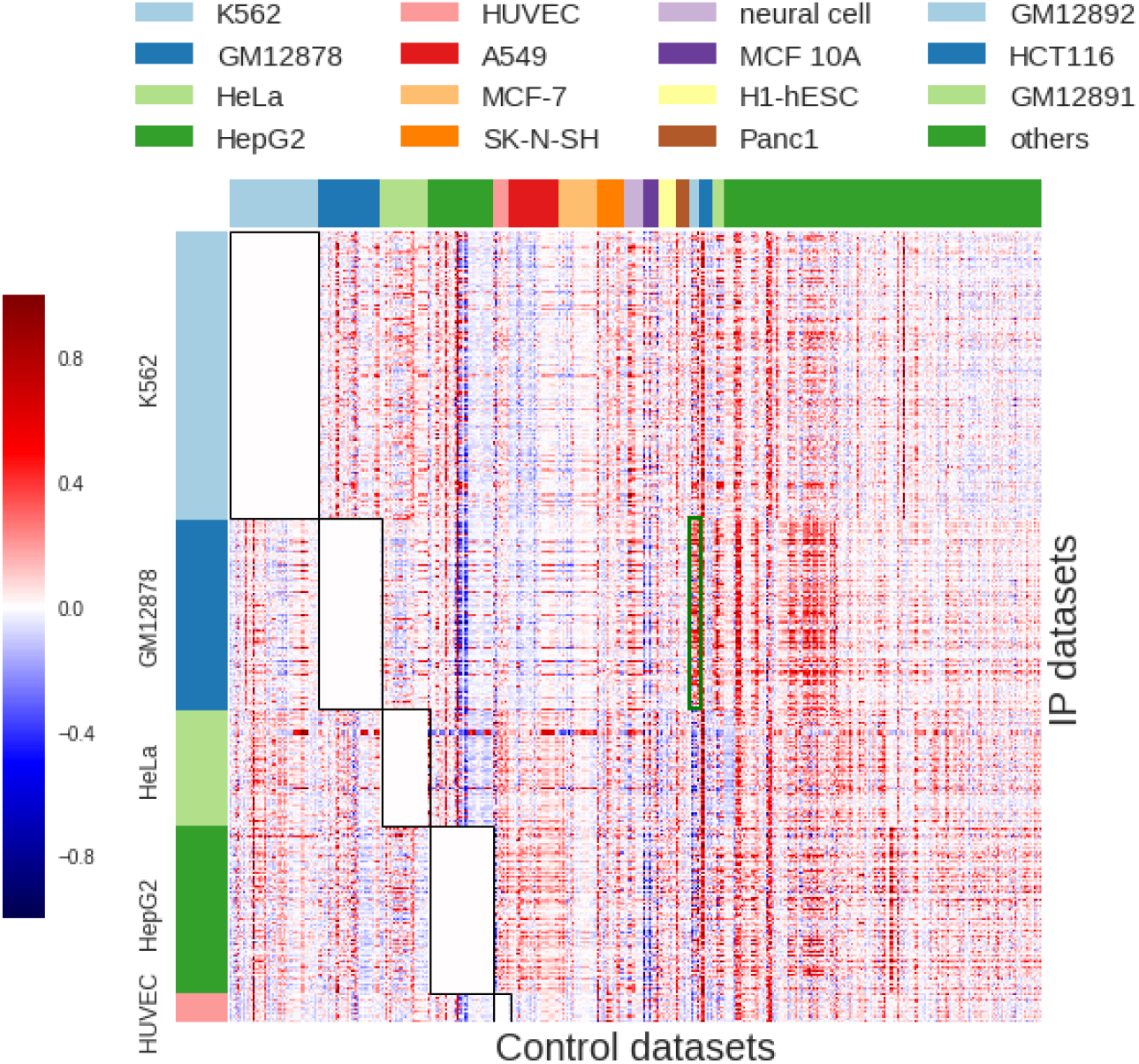
Normalized weights the AIControl model assigns to 440 ENCODE control datasets (columns) for each of the 410 IP datasets (rows) sorted by cell types (Tables S1 and S3). Control datasets from the matched cell types are excluded. The black rectangles indicate the weights of the control samples measured in the same cell line as the IP datasets .

**Figure S16.**
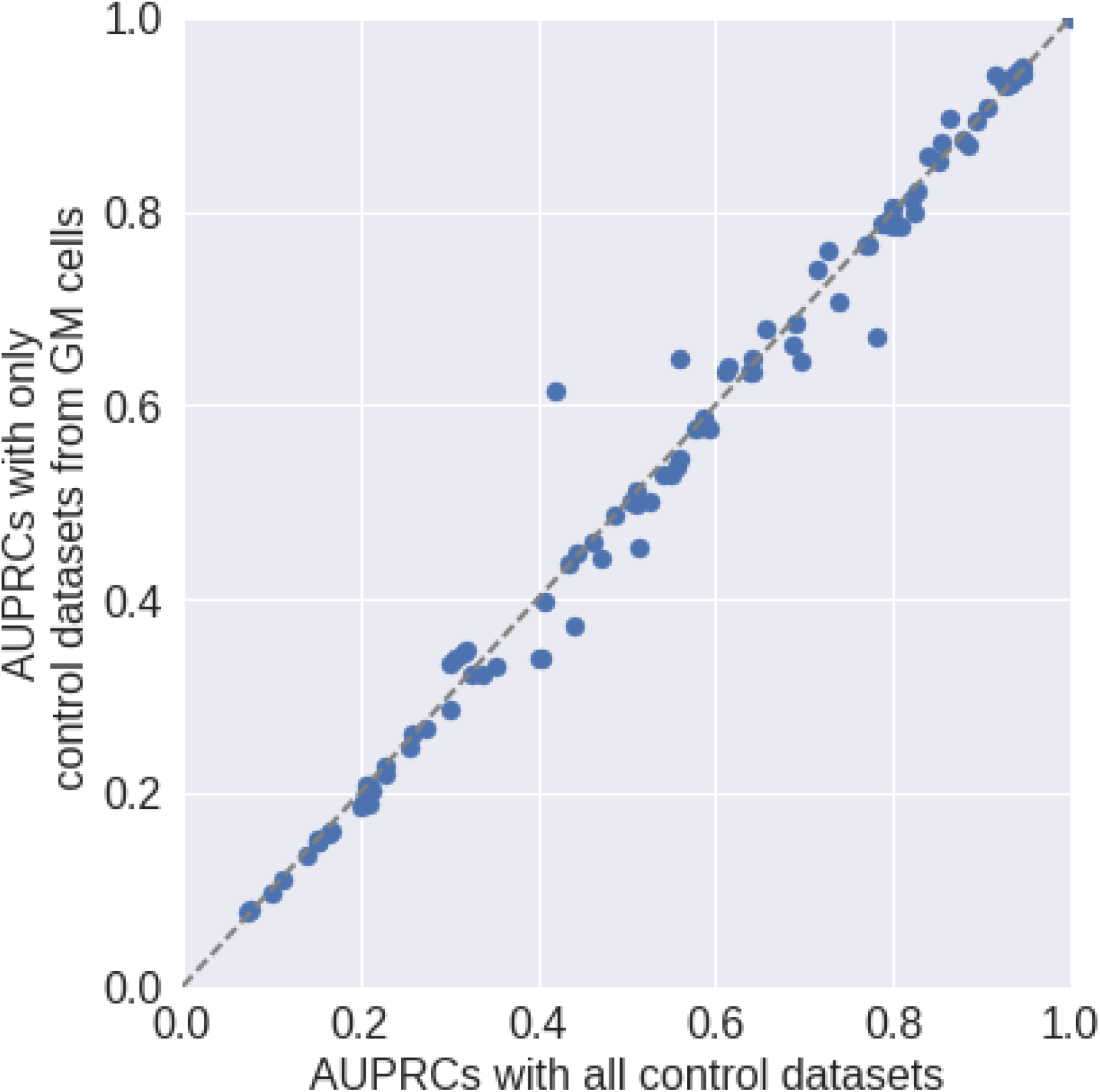
Comparison of AUPRCs from AIControl with all 440 control datasets vs. AIControl with only 93 control from the GM cell lines on the IP datasets of GM12878. Control datasets from matched experiments are removed. The performance improves with inclusion of control datasets from celltypes with abnormal karyotypes. (p-value=0.02 with Wilcoxon signed rank test)

**Figure S17.**
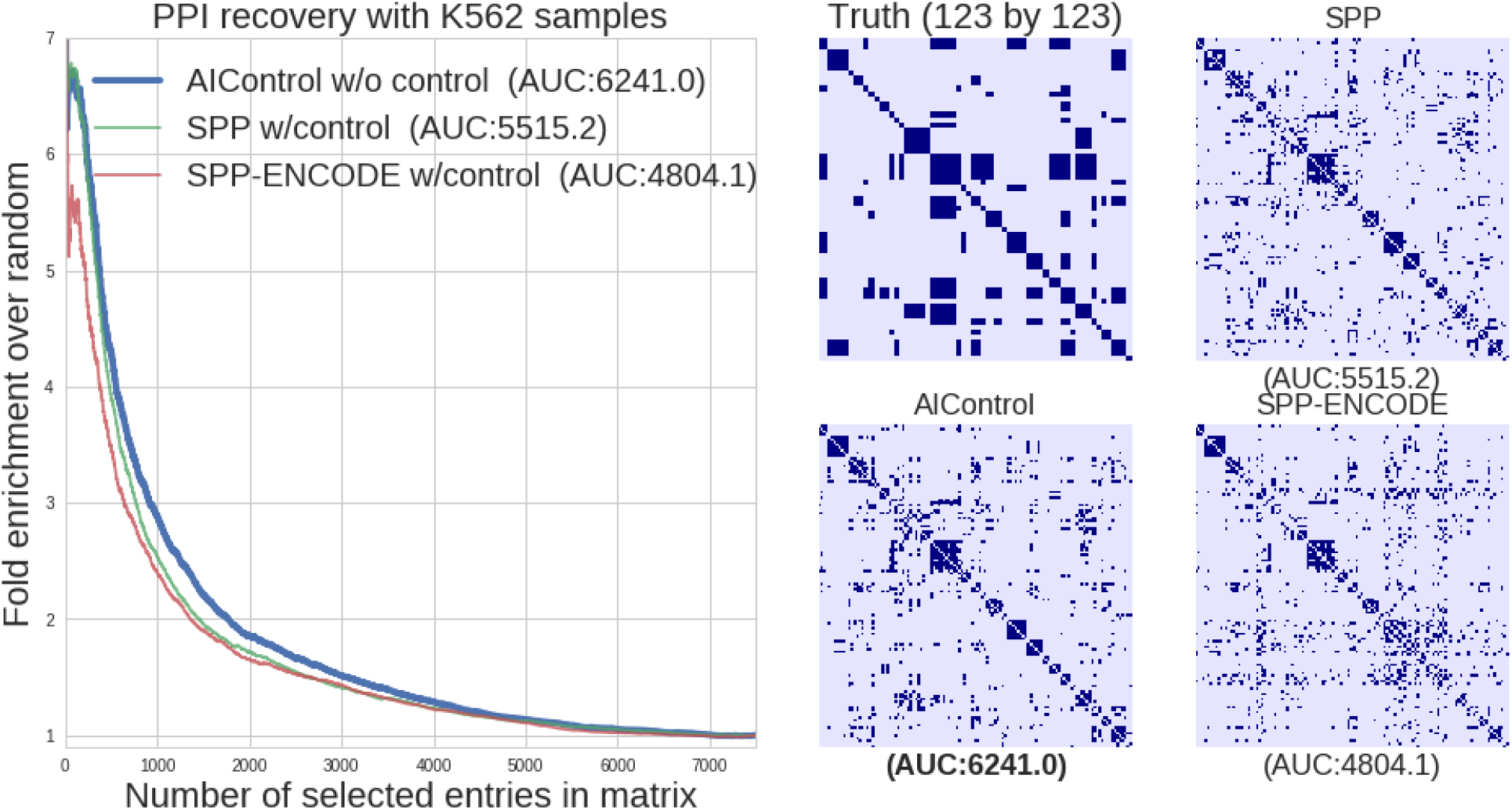
Performance of AIControl compared to other peak callers on the PPI recovery task in K562. (a) Enrichment of BioGrid-supported interactions of transcription factors in the inverse correlation networks inferred from 123 K562 IP datasets. Peak signals are obtained from: (1) AIControl w/o control, (2) SPP w/control, and (3) SPP peaks from the ENCODE database (“SPP-ENCODE”). **(b) Documented interactions among regulatory proteins (top left) and heatmaps of inverse correlation networks (rest)**. The heatmaps are binarized to show top 2269 interactions, which is equal to the number of true interactions.

**Figure S18.**
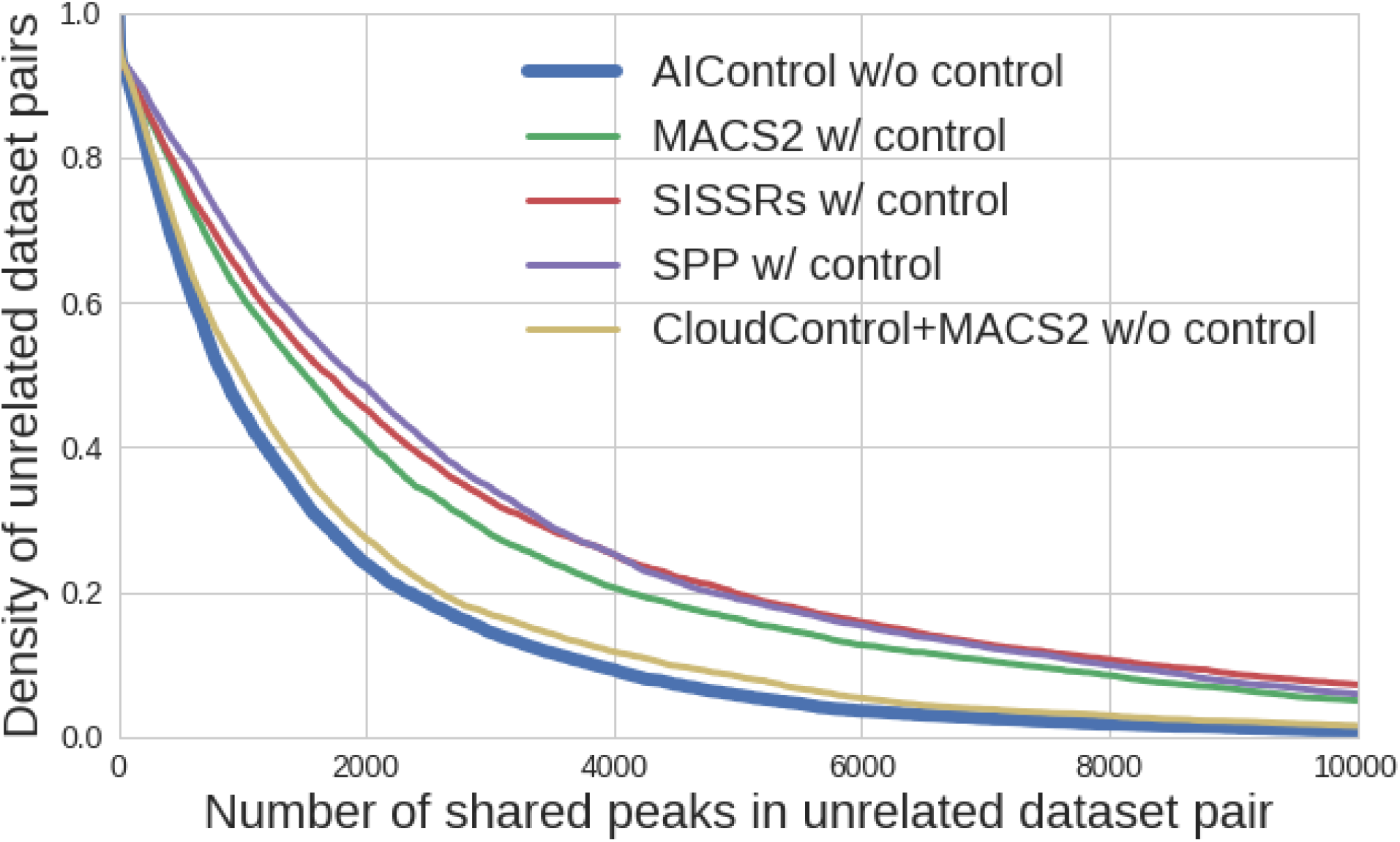
Cumulative density plot of the number of reproducible peaks for 9,310 pairs of unrelated datasets in K562 processed by five peak callers: (1) AIControl w/o control, (2) MACS2 w/control, (3) SISSRSs w/control, (4) SPP w/control, and (5) CloudControl + MACS2 w/o control. The smaller area under the curve indicates that the unrelated pairs generally have fewer shared peaks (i.e., better performance on removing `reproducible-background-signal`).

**Figure S19.**
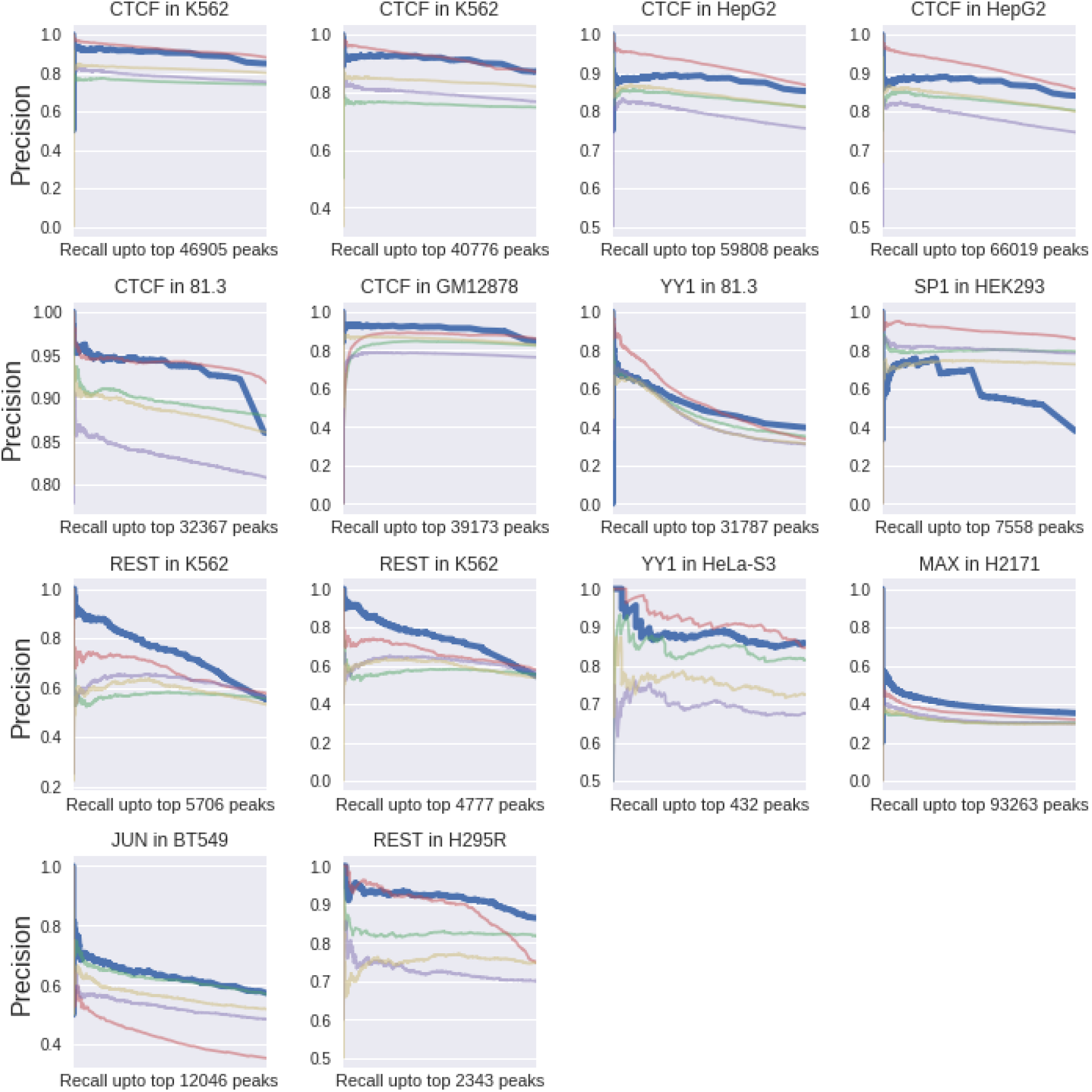
Individual precision-recall (PR) curves for predicting transcription factor binding locations using JASPAR sequence motifs as truth. All 14 external datasets are shown. The blue, green, red, purple and yellow lines indicate the performance of AIControl, MACS2, SISSRs, SPP, and Baseline (MACS2 w/o control), respectively. The AUPRC values are summarized in Figure 6.

**Figure S20.**
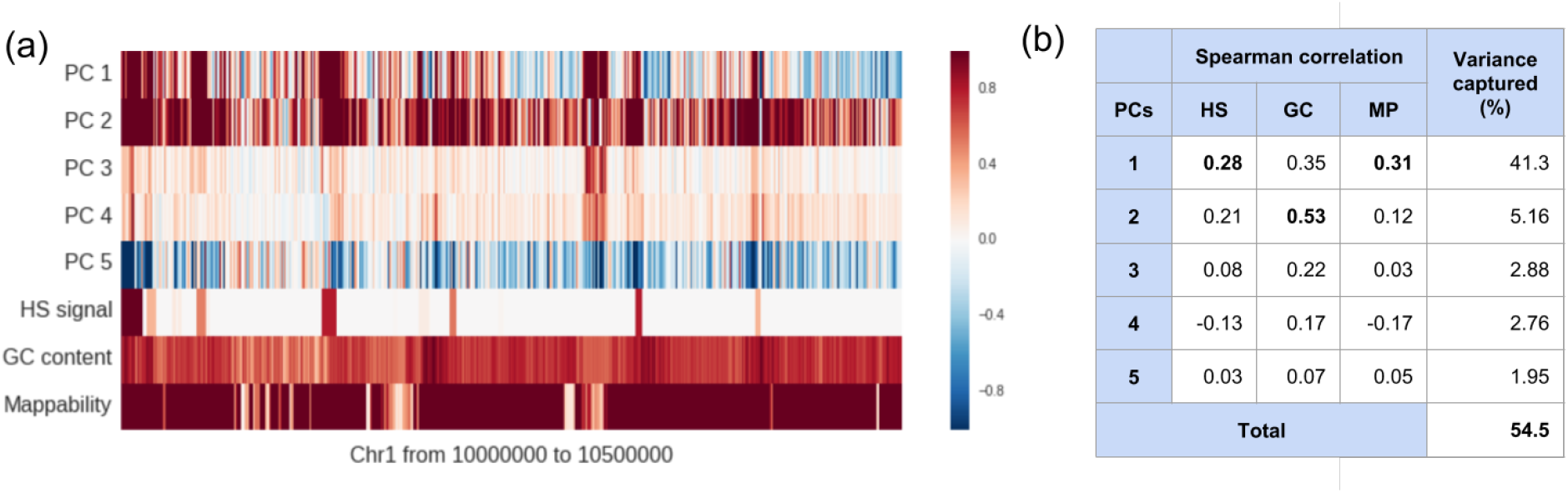
Principal component analysis of K562 control signals. (a) Normalized signals from 53 K562 control datasets projected onto 5 principal components and known bias factors on chromosome 1 from position 10,000,000 to 10,500,000. Signals are binned into 100 bp bins. (b) Spearman correlation between projected control signals and known biases: DNase 1 hypersensitivity (HS), normalized GC content (GC), and read mappability values (MP).

### Supplementary Tables

**Table S1.** Summary of all ENCODE IP datasets processed and evaluated by AIControl broken down by cell line. This summarizes the full listing of all 410 IP ChIP-seq datasets. See Supplementary Data 1 for the full list. The “Datasets” column represents how many datasets were included in the analysis as IP and control experiments, and the “transcription factors” column indicates how many unique transcription factors were included.

| Cell line | Datasets | Transcription factors |
| --- | --- | --- |
| **K562** | 149 | 40 |
| **GM12878** | 99 | 36 |
| **HepG2** | 87 | 32 |
| **HeLa-S3** | 60 | 25 |
| **HUVEC** | 15 | 5 |
| **Total** | **410** | **54** |

**Table S2.**
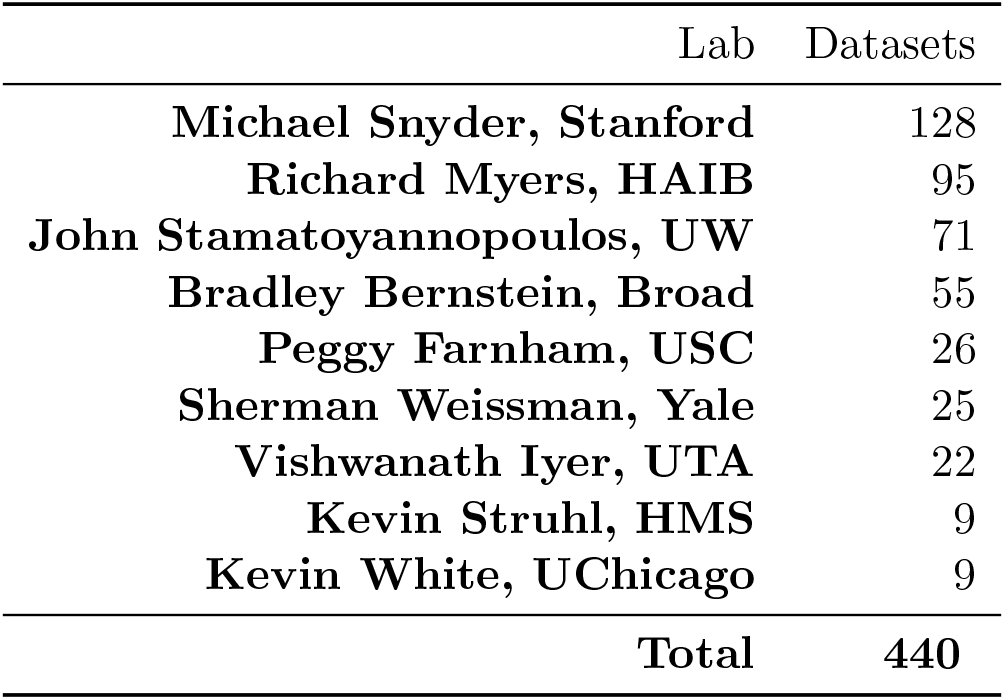
Summary of all ENCODE control datasets processed and evaluated by AIControl broken down by lab. This summarizes the full listing of all control datasets. See Supplementary Data 1 for the full list.

**Table S3.**
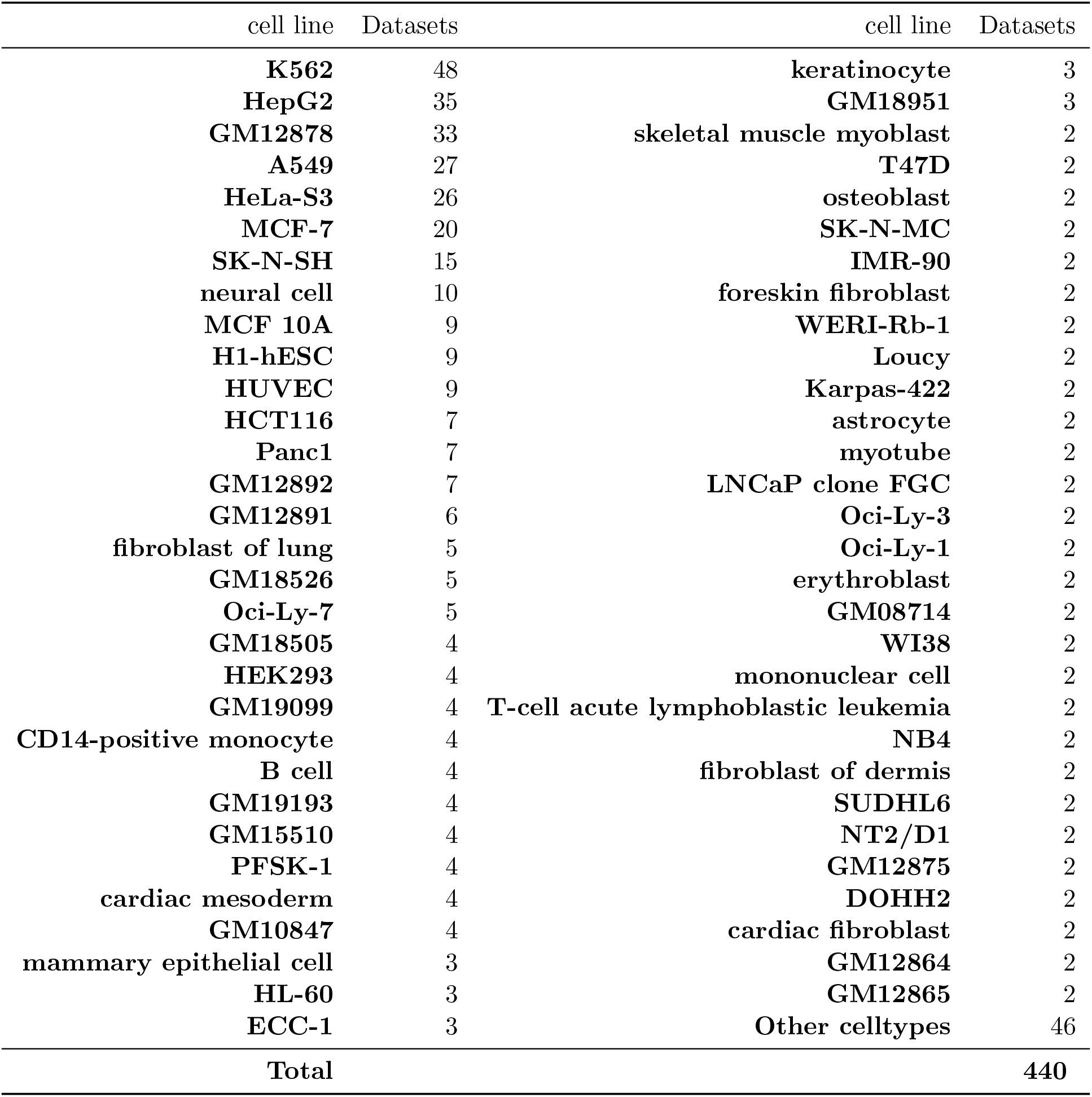
Summary of all ENCODE control datasets processed and evaluated by AIControl broken down by cell line. This summarizes the full listing of all control datasets. See Supplementary Data 1 for the full list.

| cell line | Datasets | cell line | Datasets |
| --- | --- | --- | --- |
| **K562** | 48 | **keratinocyte** | 3 |
| **HepG2** | 35 | **GM18951** | 3 |
| **GM12878** | 33 | **skeletal muscle myoblast** | 2 |
| **A549** | 27 | **T47D** | 2 |
| **HeLa-S3** | 26 | **osteoblast** | 2 |
| **MCF-7** | 20 | **SK-N-MC** | 2 |
| **SK-N-SH** | 15 | **IMR-90** | 2 |
| **neural cell** | 10 | **foreskin fibroblast** | 2 |
| **MCF 10A** | 9 | **WERI-Rb-1** | 2 |
| **H1-hESC** | 9 | **Loucy** | 2 |
| **HUVEC** | 9 | **Karpas-422** | 2 |
| **HCT116** | 7 | **astrocyte** | 2 |
| **Panc1** | 7 | **myotube** | 2 |
| **GM12892** | 7 | **LNCaP clone FGC** | 2 |
| **GM12891** | 6 | **Oci-Ly-3** | 2 |
| **fibroblast of lung** | 5 | **Oci-Ly-1** | 2 |
| **GM18526** | 5 | **erythroblast** | 2 |
| **Oci-Ly-7** | 5 | **GM08714** | 2 |
| **GM18505** | 4 | **WI38** | 2 |
| **HEK293** | 4 | **mononuclear cell** | 2 |
| **GM19099** | 4 | **T-cell acute lymphoblastic leukemia** | 2 |
| **CD14-positive monocyte** | 4 | **NB4** | 2 |
| **B cell** | 4 | **fibroblast of dermis** | 2 |
| **GM19193** | 4 | **SUDHL6** | 2 |
| **GM15510** | 4 | **NT2/D1** | 2 |
| **PFSK-1** | 4 | **GM12875** | 2 |
| **cardiac mesoderm** | 4 | **DOHH2** | 2 |
| **GM10847** | 4 | **cardiac fibroblast** | 2 |
| **mammary epithelial cell** | 3 | **GM12864** | 2 |
| **HL-60** | 3 | **GM12865** | 2 |
| **ECC-1** | 3 | **Other celltypes** | 46 |
| **Total** |  |  | **440** |

**Table S4.** A summary of the 14 external datasets that are used to analyze the generalizability of AIControl outside of ENCODE database.

| Accession | Target TF | Cell line | Matched control | Control from same cell line | Source |
| --- | --- | --- | --- | --- | --- |
| **SRR575892** | MAX | H2171 | available | none | [27] |
| **SRR941186** | REST | NCIH295R | available | none | [11] |
| **ERR011988** | CTCF | HepG2 | available | available | [38] |
| **ERR011987** | CTCF | HepG2 | available | available | [38] |
| **ERR231607** | CTCF | 81.3 | available | none | [40] |
| **ERR231612** | CTCF | GM12878 | available | available | [40] |
| **ERR231628** | YY1 | 81.3 | available | none | [40] |
| **SRR830488** | JUN | BT549 | available | none | [49] |
| **SRR332364** | YY1 | HeLa-S3 | available | available | [29] |
| **ERR637886** | SP1 | Hek293 | available | available | [46] |
| **SRX1080000** | CTCF | K562 | available | available | [37] |
| **SRX1080001** | CTCF | K562 | available | available | [37] |
| **SRX1080013** | REST | K562 | available | available | [37] |
| **SRX1080014** | REST | K562 | available | available | [37] |

**Table S5.** A table that summarizes Figure S5 by calculating the area under the water fall plots. Any performance above baseline positively contributes to the area under curve values. Alternatively, any performance below baseline negatively contributes to the values. The evaluation is performed across the whole genome. See Supplementary Data 2 for raw AUPRC values.

|  | AIControl | MACS2 | SISSRs | SPP | CloudControl |
| --- | --- | --- | --- | --- | --- |
| K562 | 37.78* | 3.65 | 25.97 | 8.53 | 12.60 |
| GM12878 | 26.93* | 7.46 | 22.15 | 14.65 | 10.71 |
| HepG2 | 33.71* | 10.20 | 26.23 | 15.69 | 12.57 |
| HeLa-S3 | 14.71* | 2.61 | 11.84 | 5.28 | 2.93 |
| HUVEC | 2.66 | -0.17 | 2.78* | -0.18 | 0.19 |

**Table S6.** A table that summarizes Figure S6 by calculating the area under the water fall plots. Any performance above baseline positively contributes to the area under curve values. Alternatively, any performance below baseline negatively contributes to the values. The evaluation is constrained in the DNase 1 hypersensitivity regions. See Supplementary Data 3 for raw AUPRC values.

|  | AIControl | MACS2 | SISSRs | SPP | CloudControl |
| --- | --- | --- | --- | --- | --- |
| K562 | 26.48* | 2.09 | 20.84 | 4.68 | 9.58 |
| GM12878 | 18.03* | 3.42 | 14.17 | 7.75 | 6.87 |
| HepG2 | 19.09* | 1.88 | 13.46 | 5.82 | 6.48 |
| HeLa-S3 | 8.99* | 0.11 | 6.44 | 1.93 | 1.99 |
| HUVEC | 1.90* | -0.04 | 1.82 | -0.08 | 0.15 |

**Table S7.**
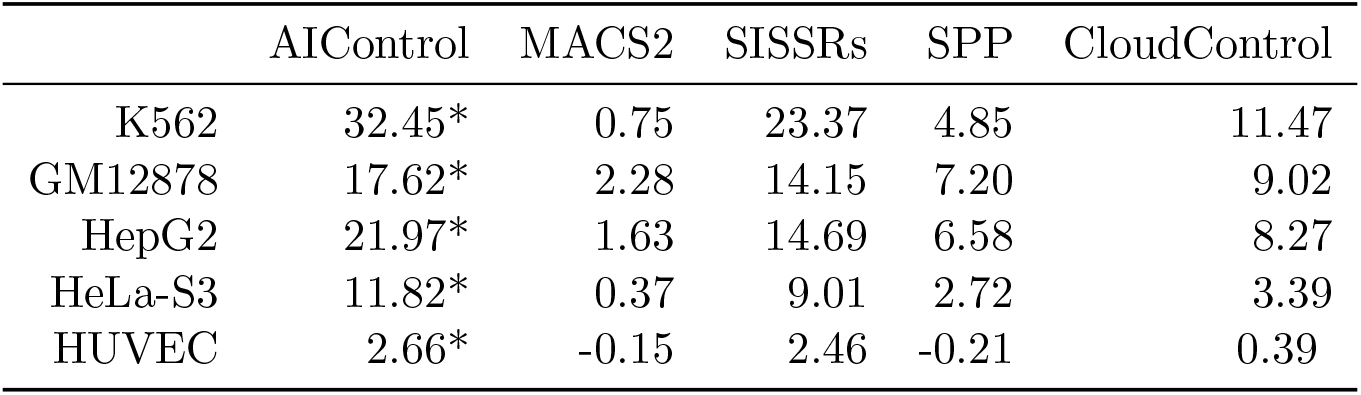
A table that summarizes Figure S7 by calculating the area under the water fall plots. Any performance above baseline positively contributes to the area under curve values. Alternatively, any performance below baseline negatively contributes to the values. The evaluation is constrained in 5 kilo-base up and down stream of the start of protein coding genes. See Supplementary Data 4 for raw AUPRC values.

**Table S8.** A table that summarizes Figure S8 by calculating the area under the water fall plots. Any performance above baseline positively contributes to the area under curve values. Alternatively, any performance below baseline negatively contributes to the values. The evaluation is constrained in regions of the hg38 genome with more than 50% GC content. See Supplementary Data 5 for raw AUPRC values.

|  | AIControl | MACS2 | SISSRs | SPP | CloudControl |
| --- | --- | --- | --- | --- | --- |
| K562 | 27.37* | -0.65 | 15.39 | 2.20 | 8.47 |
| GM12878 | 17.00* | 3.17 | 10.42 | 5.51 | 9.40 |
| HepG2 | 18.53* | -0.62 | 12.17 | 2.98 | 5.35 |
| HeLa-S3 | 9.91* | -0.06 | 6.62 | 1.19 | 2.83 |
| HUVEC | 2.33* | -0.09 | 2.29 | -0.20 | 0.26 |

**Table S9.**
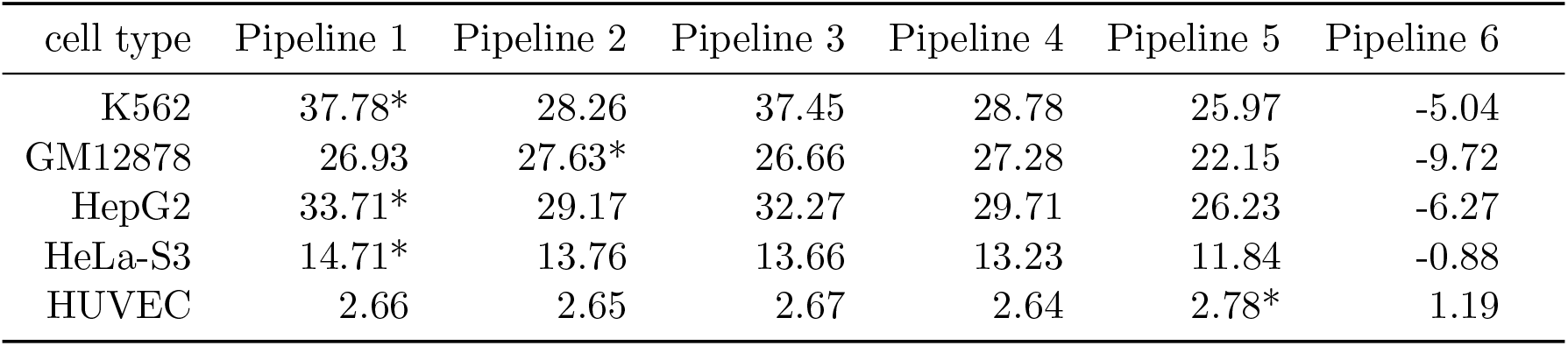
A table that summarizes Figure 4 by calculating the area under the water fall plots. Any performance above baseline positively contributes to the area under curve values. Alternatively any performance below baseline negatively contributes to the values. The evaluation is constrained in the DNase 1 hypersensitivity regions. Following pipelines were compared: (1) AIControl with all 440 control datasets except for matched controls, (2) AIControl without control datasets from the same cell type as the IP dataset, (3) AIControl without control datasets from the same lab as the IP dataset, (4) AIControl without control datasets from the same lab or the same cell type as the IP dataset, (5) SISSRs with matched control datasets, and (6) SISSRs without matched control datasets. See Supplementary Data 2 for raw AUPRC values.

**Table S10.** A list of true PPIs in K562 that are supported by the BioGrid Database, and the number of edges recovered by the six peak callers: (A) AIControl w/o control, (B) MACS2 w/ control, (C) SISSRs w/ control, (D) SPP w/ control, (E) CloudControl+MACS2 w/o control, and (F) MACS2 w/o control. An edge is considered as ‘‘recovered”, if it is in the 3,583 most significant predictions. The threshold value (3,583) is equal to the number of positive entries in the truth matrix.

| PPI |  | # of edges recovered |  |  |  |  |  | PPI |  | # of edges recovered |  |  |  |  |  |
| --- | --- | --- | --- | --- | --- | --- | --- | --- | --- | --- | --- | --- | --- | --- | --- |
| TF1 | TF2 | A | B | C | D | E | F | TF1 | TF2 | A | B | C | D | E | F |
| CTCF | ZBTB33 | 1 | 1 | 3 | 2 | 1 | 4 | NFYA | NFYB | 6 | 7 | 6 | 6 | 6 | 6 |
| NFYB | TBP | 4 | 1 | 1 | 2 | 3 | 1 | JUND | MEF2A | 1 | 0 | 0 | 0 | 1 | 0 |
| MAX | USF1 | 10 | 9 | 9 | 9 | 10 | 9 | JUND | YY1 | 2 | 3 | 1 | 3 | 2 | 2 |
| CEBPB | CREB1 | 1 | 1 | 1 | 1 | 1 | 1 | SP1 | TBP | 6 | 2 | 8 | 5 | 5 | 4 |
| JUN | STAT1 | 24 | 13 | 18 | 15 | 15 | 17 | CREB1 | JUN | 4 | 7 | 9 | 10 | 12 | 8 |
| E2F6 | MAX | 21 | 16 | 15 | 16 | 18 | 16 | SP1 | SRF | 2 | 4 | 2 | 4 | 4 | 4 |
| REST | SP1 | 3 | 5 | 8 | 6 | 8 | 9 | SP1 | SREBF1 | 2 | 5 | 4 | 1 | 0 | 5 |
| SREBF1 | USF1 | 0 | 0 | 0 | 0 | 0 | 0 | FOSL1 | JUN | 19 | 16 | 18 | 18 | 17 | 17 |
| ESRRA | SP1 | 4 | 4 | 6 | 6 | 7 | 6 | ELF1 | NFYA | 1 | 1 | 0 | 1 | 0 | 0 |
| CEBPB | JUN | 10 | 2 | 6 | 4 | 7 | 4 | FOS | JUN | 28 | 23 | 26 | 24 | 26 | 22 |
| JUN | TBP | 1 | 1 | 2 | 1 | 1 | 1 | EGR1 | SP1 | 5 | 2 | 3 | 3 | 3 | 3 |
| ELK1 | SRF | 4 | 3 | 3 | 3 | 4 | 3 | FOS | JUND | 0 | 0 | 0 | 0 | 0 | 0 |
| CEBPB | SRF | 0 | 0 | 0 | 0 | 0 | 0 | FOS | MAFF | 0 | 0 | 0 | 0 | 0 | 0 |
| FOS | USF2 | 5 | 5 | 5 | 4 | 5 | 5 | CEBPB | SP1 | 1 | 0 | 2 | 0 | 1 | 0 |
| MAFF | NRF1 | 0 | 1 | 0 | 0 | 1 | 0 | SP1 | YY1 | 2 | 3 | 3 | 1 | 3 | 2 |
| CEBPB | NRF1 | 0 | 1 | 1 | 0 | 0 | 0 | JUN | MEF2A | 7 | 4 | 3 | 8 | 6 | 6 |
| FOS | TBP | 0 | 1 | 0 | 1 | 0 | 1 | IRF1 | STAT1 | 33 | 16 | 20 | 25 | 23 | 20 |
| MEF2A | TBP | 0 | 0 | 0 | 0 | 0 | 0 | JUN | JUND | 15 | 16 | 15 | 14 | 15 | 14 |
| CREB1 | YY1 | 2 | 4 | 5 | 5 | 2 | 3 | FOS | STAT1 | 3 | 0 | 0 | 0 | 0 | 1 |
| NFYB | SP1 | 2 | 5 | 1 | 4 | 4 | 4 | NFYA | SRF | 0 | 0 | 0 | 1 | 0 | 0 |
| CREB1 | JUND | 4 | 3 | 4 | 3 | 4 | 3 | FOS | USF1 | 1 | 0 | 0 | 1 | 1 | 0 |
| CEBPB | RUNX1 | 0 | 0 | 0 | 0 | 0 | 0 | JUN | RUNX1 | 0 | 0 | 2 | 0 | 0 | 0 |
| IRF1 | IRF2 | 10 | 9 | 8 | 10 | 9 | 7 | FOSL1 | USF1 | 1 | 2 | 1 | 1 | 1 | 1 |
| CEBPB | EGR1 | 0 | 0 | 0 | 1 | 1 | 0 | JUN | MAX | 12 | 6 | 7 | 6 | 9 | 5 |
| EGR1 | JUND | 2 | 1 | 2 | 1 | 2 | 1 | ELF1 | SP1 | 2 | 4 | 3 | 3 | 2 | 2 |
| GABPA | YY1 | 3 | 2 | 1 | 2 | 3 | 3 | ELF1 | RUNX1 | 0 | 0 | 3 | 0 | 0 | 0 |
| GATA2 | JUN | 20 | 6 | 10 | 10 | 13 | 10 | FOS | RUNX1 | 0 | 0 | 0 | 0 | 0 | 0 |
| USF1 | USF2 | 3 | 3 | 3 | 3 | 3 | 1 | NFYA | SP1 | 6 | 5 | 4 | 5 | 4 | 5 |
| CREB1 | NFYA | 2 | 2 | 1 | 3 | 2 | 2 | SREBF1 | YY1 | 3 | 2 | 1 | 0 | 2 | 2 |
| MAX | TBP | 6 | 6 | 5 | 4 | 6 | 4 | GABPA | SP1 | 1 | 2 | 3 | 2 | 1 | 2 |
| IRF1 | SP2 | 4 | 2 | 1 | 3 | 6 | 3 | JUN | NFYA | 0 | 0 | 0 | 0 | 0 | 0 |
| MAX | YY1 | 12 | 7 | 7 | 7 | 8 | 8 | CTCF | YY1 | 7 | 7 | 3 | 11 | 7 | 4 |
| REST | TBP | 3 | 2 | 2 | 2 | 2 | 2 | JUN | SP1 | 2 | 0 | 7 | 2 | 3 | 2 |

**Table S11.**
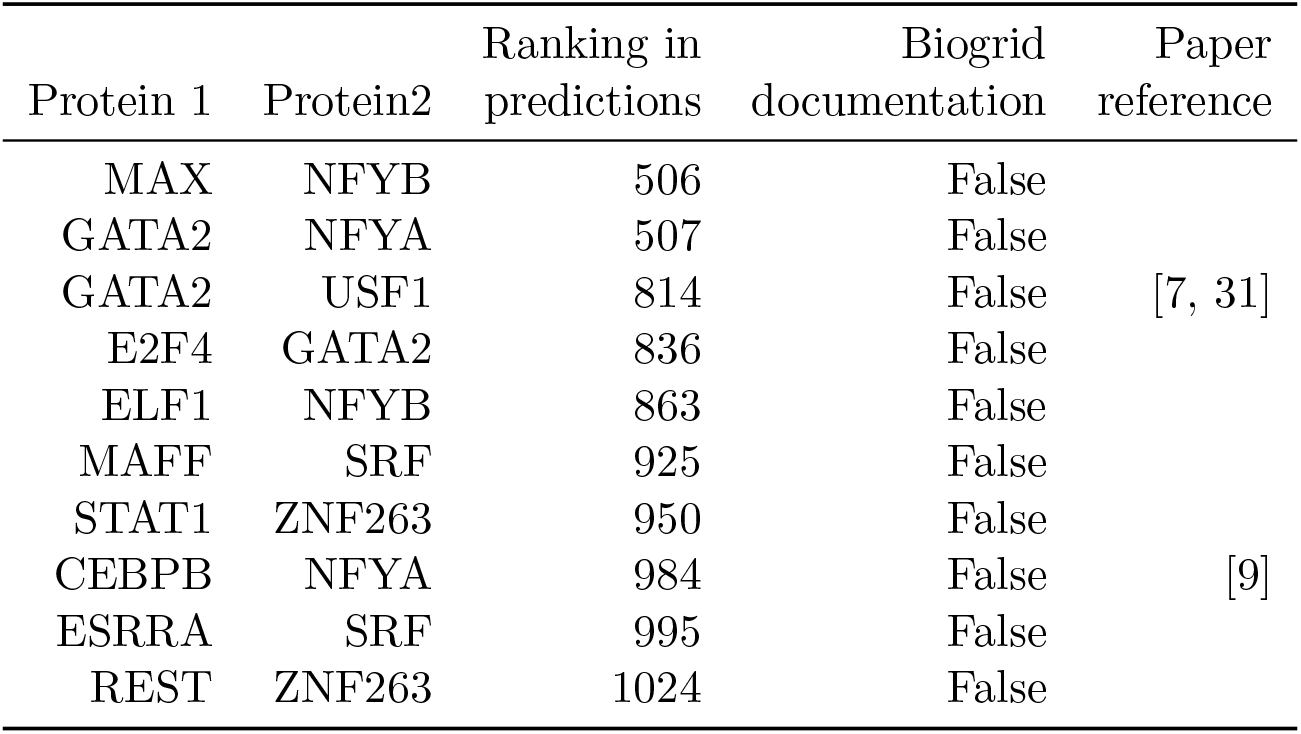
A list of top 10 PPIs that AIControl uniquely ranked in the 3,583 most significant interactions. 3,583 is the number of true positives.

## Supporting information

SupplementaryData.zip

